# The MarR-type regulator MalR is involved in stress-responsive cell envelope remodeling in *Corynebacterium glutamicum*

**DOI:** 10.1101/544056

**Authors:** Max Hünnefeld, Marcus Persicke, Jörn Kalinowski, Julia Frunzke

## Abstract

1

It is the enormous adaptive capacity of microorganisms, which is key to their competitive success in nature, but also challenges antibiotic treatment of human diseases. To deal with a diverse set of stresses, bacteria are able to reprogram gene expression using a wide variety of transcription factors. Here, we focused on the MarR-type regulator MalR conserved in the *Corynebacterineae*, including the prominent pathogens *Corynebacterium diphtheriae* and *Mycobacterium tuberculosis*. In several corynebacterial species, the *malR* gene forms an operon with a gene encoding a universal stress protein (*uspA*). Chromatin-affinity purification and sequencing (ChAP-Seq) analysis revealed that MalR binds more than 60 target promoters in the *C. glutamicum* genome as well as in the large cryptic prophage CGP3. Overproduction of MalR caused severe growth defects and an elongated cell morphology. ChAP-Seq data combined with a global transcriptome analysis of the *malR* overexpression strain emphasized a central role of MalR in cell envelope remodeling in response to environmental stresses. Prominent MalR targets are for example involved in peptidoglycan biosynthesis and synthesis of branched-chain fatty acids. Phenotypic microarrays suggest an altered sensitivity of a Δ*malR* mutant towards several β-lactam antibiotics. We furthermore revealed MalR as a repressor of several prophage genes suggesting that MalR may be involved in the control of stress-responsive induction of the large CGP3 element. In conclusion, our results emphasize MalR as a regulator involved in stress-responsive remodeling of the cell envelope of *C. glutamicum* and suggest a link between cell envelope stress and the control of phage gene expression.

**Importance:** Bacteria live in changing environments that force the cells to be highly adaptive. The cell envelope represents both, a barrier against harsh external conditions and an interaction interface. The dynamic remodeling of the cell envelope as a response towards, e.g. antibiotic treatment represents a major challenge in the treatment of diseases. Members of the MarR family of regulators are known to contribute to an adaptation of bacterial cells towards antibiotic stress. However, our knowledge on this adaptive response was so far restricted to a small number of well-described target genes. In this study, we performed a genome-wide profiling of DNA-binding of the MarR-type regulator MalR of *C. glutamicum*, which is conserved in several coryne- and mycobacterial species. By binding to more than 60 different target promoters, MalR is shaping a global reprogramming of gene expression conferring a remodeling of the cell envelope in response to stress.

## 2 Introduction

In almost every natural habitat a high number of microbial species coexist and compete for space and nutrients. In this competitive scenario, the exposure to bacteriostatic or bactericidal compounds (antibiotics) represents a routine challenge which bacteria are facing in various natural environments and particularly during infection of a specific host (1–3). A prominent family of transcription factors involved in the reprogramming of gene expression in response to stress conditions is the MarR-type family (4, 5). Already decades ago, clinical isolates of *Escherichia coli* displaying a <u>m</u>ultiple <u>a</u>ntibiotic <u>r</u>esistance phenotype where found to carry mutations in the *marR* locus (6) and subsequently drew considerable attention to this ubiquitously found class of regulators. Following studies then showed that *E. coli* MarR is a transcriptional repressor of genes conferring resistance towards different antibiotics, organic solvents and lipophilic, mainly phenolic compounds (7). In further studies it was shown that MarR-type regulators are widely distributed among bacteria and archaea, likely representing an ancient regulator family which emerged before the evolutionary split of these domains more than three billion years ago (8, 9). Overall, the regulatory responses modulated by MarR-type regulators were grouped into three general categories (4), including (i) environmental stress responses (*e.g.* triggered by antibiotics) (10–12), (ii) regulation of virulence genes (13, 14) and (iii) degradation of lipophilic (often aromatic) compounds (15, 16). The DNA-binding domain of MarR-family regulators is typically comprised of a winged helix-turn-helix domain recognizing palindromes or inverted repeats (17). In the classical scenario, the dissociation of the MarR dimer from its genetic target is triggered by ligand binding (*e.g.* antibiotics, salicylates, lipophilic compounds (18)), but also examples exist where the binding of ligands fosters the association to DNA targets (15, 19).

The suborder of the *Corynebacterineae* covers several prominent pathogenic species, such as *Corynebacterium diphtheriae*, *Mycobacterium tuberculosis*, and *Mycobacterium leprae* causing millions of deaths every year. Species of this suborder share a very similar and unique cell wall composition hampering antibiotic treatment (20–24). In addition to the peptidoglycan, cells are surrounded by an arabinogalactan zone topped by a lipid bilayer composed of long-chain α-alkyl, β-hydroxy fatty acids - the mycolic acids (25).

In this study, we have characterized the function of the MarR-type regulator MalR (Cg3315) of the non-pathogenic, Gram-positive model organism *C. glutamicum* (26), which – in total - harbors nine MarR-type regulators (27). Further, the genome of *C. glutamicum* contains a large prophage element (CGP3) which was shown to be inducible by the cellular SOS response (28) or excises spontaneously in a small fraction of wild type cells (29, 30).

*C. glutamicum* MalR was previously published as a repressor of the *malE* gene, encoding the malic enzyme (31). Here, we performed a genome-wide profiling of MalR targets by combining ChAP-Seq and a comparative transcriptomics approach. As revealed by phenotypic microarrays, a mutant lacking the *malR* gene displayed an impaired resistance towards different β-lactam antibiotics. The majority of former studies focused on a very distinct operon or small regulon controlled by MarR-type regulators. The present study provides - for the first time - a comprehensive insight into the complex regulon of MalR, which is involved in the remodeling of the cell envelope in response to stress conditions. Interestingly, our data also suggest a role of MalR in the control of the large prophage CGP3 and thereby demonstrate a complex regulatory interaction between the host and horizontally-acquired elements.

## 3 Results

### The MarR-type regulator MalR is conserved in Corynebacteria and Mycobacteria

The MarR-type regulator MalR (Cg3315) was previously described as a repressor of the malic enzyme gene in *C. glutamicum* (31). Sequence analysis revealed that MalR is conserved in several coryne- and mycobacterial species also including the prominent pathogens *C. diphtheriae* (57 % sequence identity) and *M. leprae* (40 % sequence identity). Simulated secondary structures of MalR using Phyre^2^ disclosed a high similarity to the secondary structure of MarR from *E. coli* consisting of six α-helices surrounding two β-sheets (32), although the amino acid sequence identity is only 22% (Fig. S1).

In the genome of *C. glutamicum* ATCC 13032, *malR* is organized in an operon with a gene encoding a universal stress protein (*uspA*) and divergently located to a small hypothetical protein followed by an operon coding for a penicillin-binding protein and two putative membrane proteins (Fig. 1) (33). This genomic ensemble emphasizes a role of MalR in global stress responses and potentially cell envelope-related functions. The genome-wide analysis of MalR target genes and its physiological impact is the aim of the present study.

**Figure 1:**
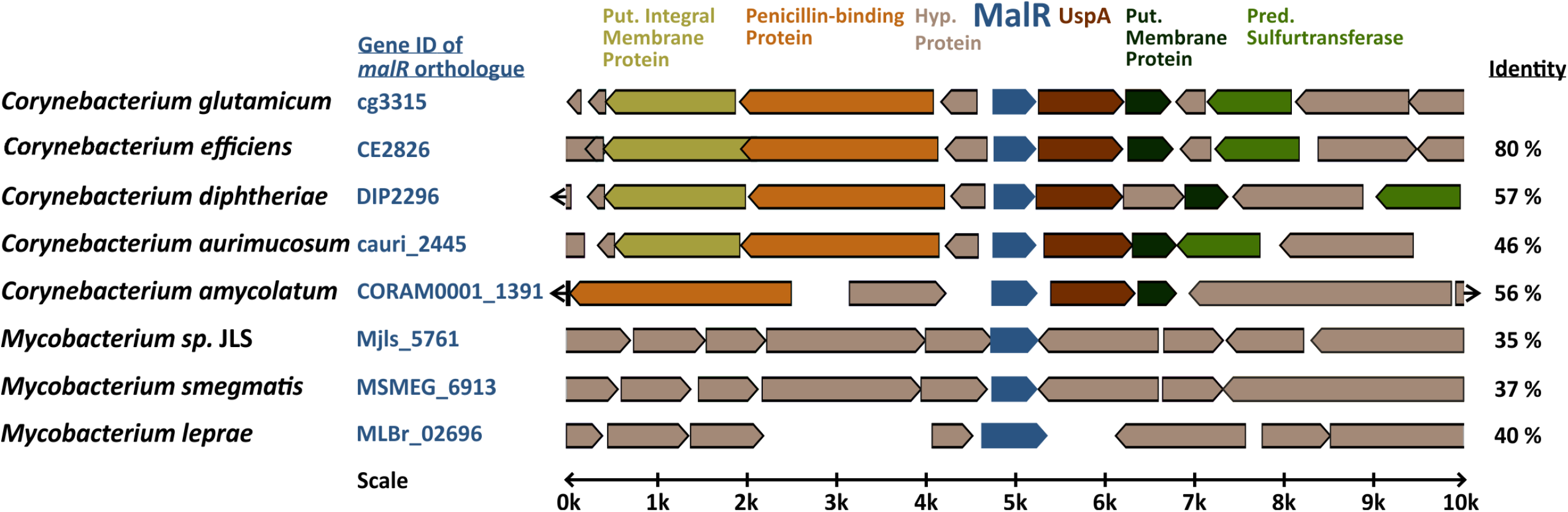
Genomic organization of the *malR* gene in coryne- and mycobacterial species. Amino acid sequence identity to the *C. glutamicum* MalR ortholog is given in the right column. The genomic context of *C. glutamicum malR* was extracted from microbesonline (http://microbesonline.org).

### Genome-wide profiling of MalR target genes

In order to identify target genes of MalR, ChAP-Seq analysis was performed and selected target promoters were subsequently verified using electrophoretic mobility shift assays (EMSA) (Fig. 2). To produce MalR at physiological levels, a gene fusion was integrated at the *malR* locus into the *C. glutamicum* ATCC 13032 chromosome, encoding a C-terminally strep-tagged variant of MalR. Cells were grown in CGXII minimal medium with 2 % glucose and harvested in the mid-exponential phase. The sequencing of DNA bound to MalR under the chosen conditions revealed 66 binding regions in total (Fig. 2 A; Tab. S1) Remarkably, 13 target regions of MalR were found inside the cryptic prophage element CGP3 showing a local maximum in the region cg1895-cg1950 (8 peaks). Besides several genes of unknown function, MalR bound to promoter regions of genes involved in cell envelope biosynthesis, including *embC* (34), the *murAB* operon (35) and *ipsA* (36). It further associates to promoter regions of genes encoding for proteins involved in transport mechanisms, such as *oppA*, cg1454, cg2256 and cg2340.

**Figure 2:**
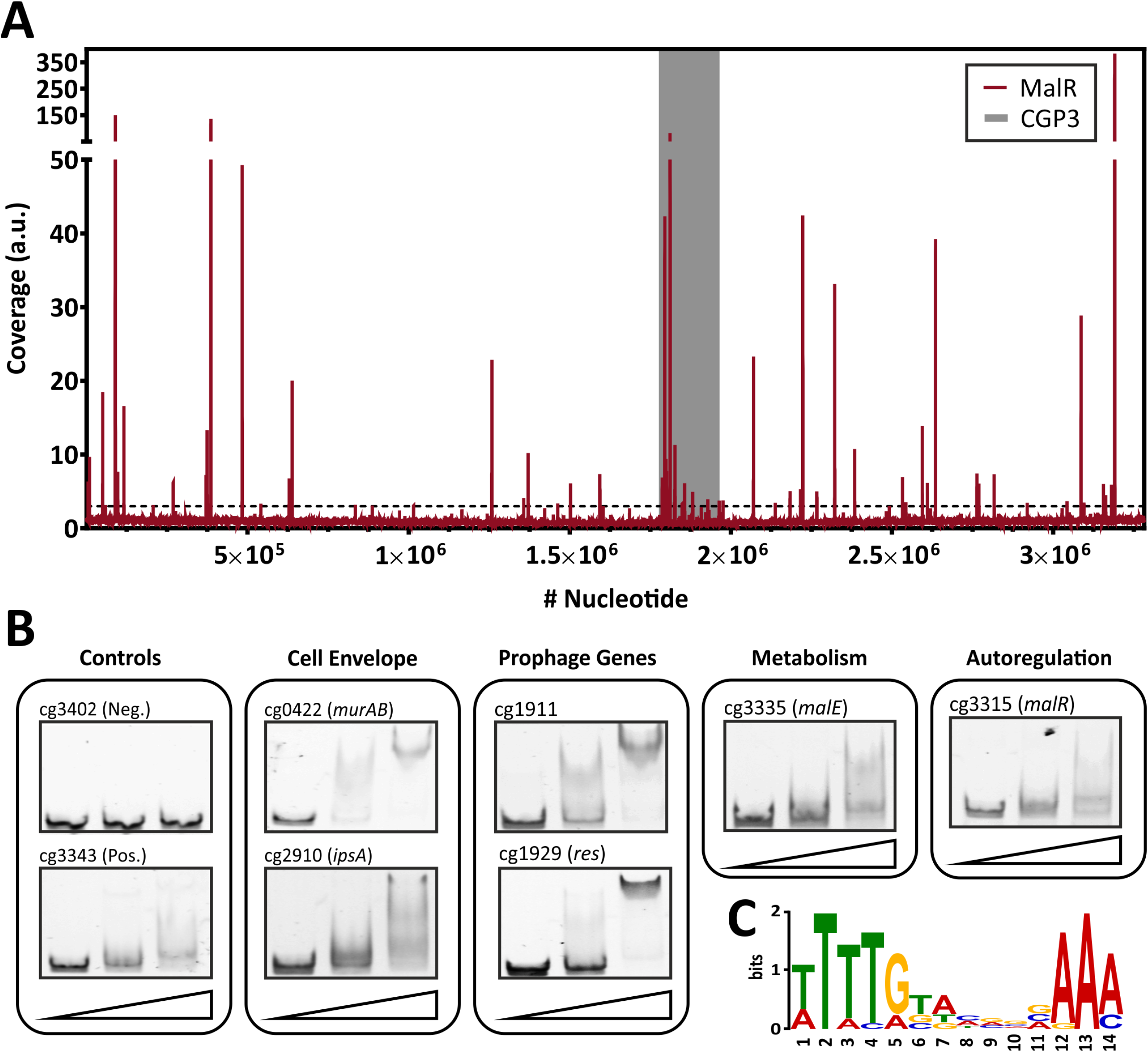
Genome-wide binding profile and *in vitro* verification of binding sites of MalR. **(A)** ChAP-Seq experiments revealed 66 distinct binding peaks of MalR in the genome of *C. glutamicum* ATCC 13032. For ChAP-Seq analysis, a strain containing a genomically encoded Strep-tagged variant of MalR was grown in CGXII medium containing 2% (w/v) glucose for 5 hours and treated as described previously (75). The coverage of MalR-bound DNA regions (red) was normalized using a rolling mean with a step size of 10 bp and a window size of 50 bp. The grey bar marks the CGP3 prophage region inside of the genome of *C. glutamicum*. Overall, 66 peaks with coverages higher than 3-fold mean coverage were detected. **(B)** Electrophoretic mobility shift assays (EMSAs) were performed to verify binding of MalR to promoter regions identified via ChAP-Seq. Therefore, 100 bp DNA fragments (50 bp up- and downstream the peak maximum) were incubated without protein (first lane), with 3 molar excess (228 nM, second lane) and 10 molar excess of purified MalR (760 nM, third lane). A complete overview on all tested fragment is shown in Fig. S2 A. **(C)** DNA sequences of 16 binding sites that were verified using EMSAs were used to deduce a possible binding motif of MalR with the online tool MEME-ChIP (48). A motif based on all peaks obtained by ChAP-Seq, as well as the distribution of the motifs within the uploaded sequences, are shown in Fig. S3.

Conspicuously, 16 peaks were detected in promoter region of genes coding for (putative) secreted proteins. Consistent with the report of Krause and coworkers, also binding to the promoter region of *malE*, was confirmed by our study (31). Furthermore, a significant binding peak was observed in the own promoter region of the *malR*-*uspA* operon indicating an auto regulation of *malR* expression. In summary, this ChAP-Seq analysis revealed a global role of MalR in the regulation of genes involved in cell envelope-related functions and suggested a regulatory interaction of MalR with the large prophage CGP3.

To validate the obtained binding profile of MalR, EMSAs were performed using different promoter regions identified by ChAP-seq analysis (Fig. 2 B). Except one potential target promoter (cg2962), every tested candidate could be verified using this *in vitro* approach (Fig. S2). As a negative control, the promoter region of cg3402 (a putative copper chaperone) was used. Here, no shift was detectable. In vitro, different migration patterns were observed for the tested MalR targets, which likely reflect differences in binding affinities and/or the presence of multiple DNA motifs. Furthermore, in some cases additional factors may contribute to in vivo MalR-DNA association (e.g. in the case of cg2962).

Using the 66 MalR peak sequences, a putative binding motif of MalR was deduced using the online tool MEME-ChIP (37). This tool predicted a very AT-rich palindromic binding motif found in all peaks (motif and distribution in Fig. S3), which is very similar to the motif found in MalR targets verified with EMSAs (Fig. 2 C).

### Overproduction of MalR causes severe growth defects

In a next step, we compared the growth of the *C. glutamicum* wild type with a *malR* deletion strain (Δ*malR*) and a strain overexpressing the *malR* gene under control of the IPTG-inducible P_*tac*_ promoter (Fig. 3). In fact, overexpression of *malR* caused a severe growth defect of *C. glutamicum* grown on CGXII minimal medium with 2 % glucose (Fig. 3 A), whereas the deletion mutant had only a minor impact on the growth rate compared to the wild type strain under the tested conditions (Fig. 3 B).

**Figure 3:**
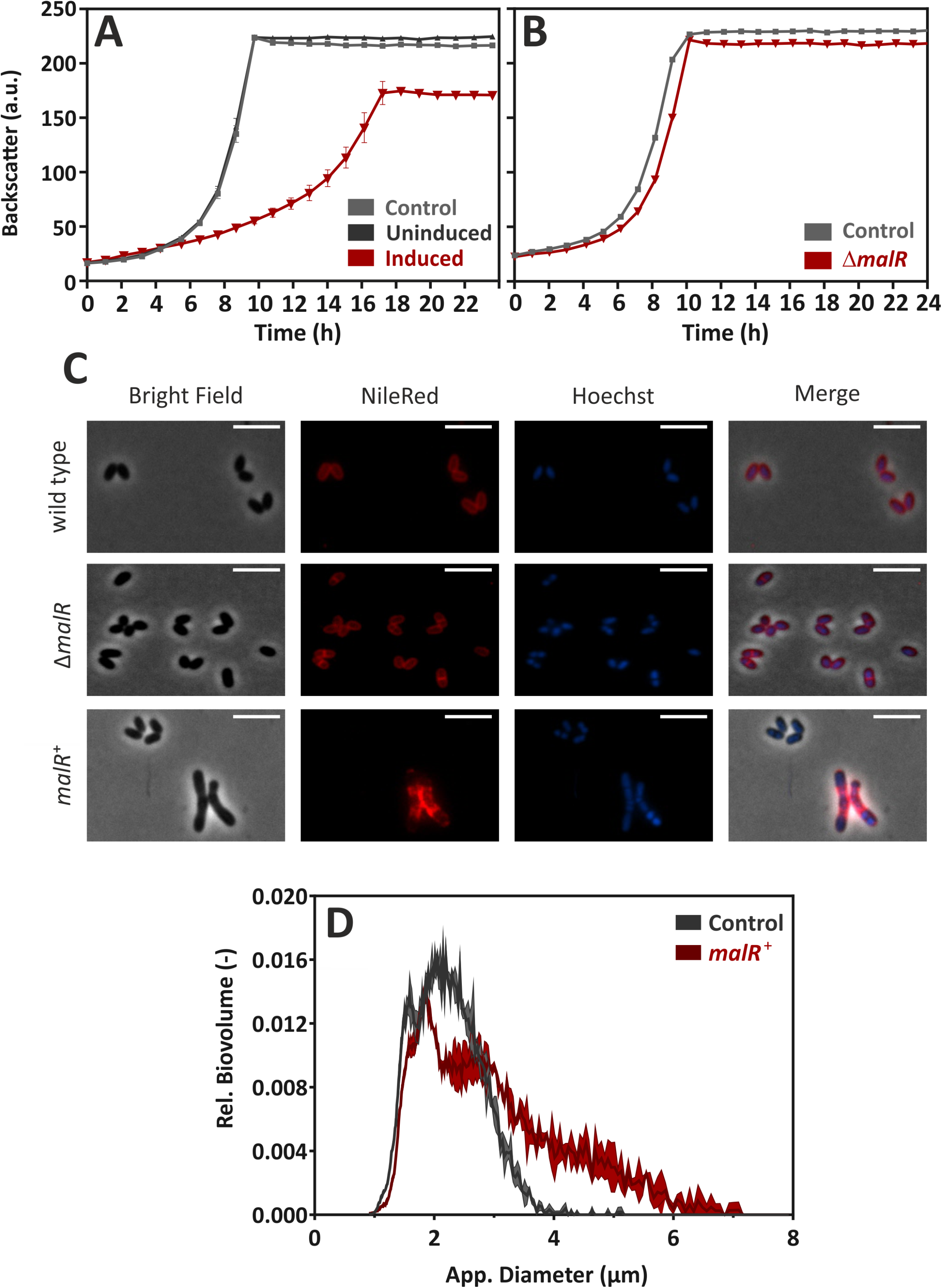
MalR overproduction causes severe growth defects of *C. glutamicum*. **(A)** Comparative growth experiment of *C. glutamicum* ATCC 13032 carrying the empty vector pAN6 and strain *C. glutamicum*/pAN6-malR overexpressing the *malR* gene. Cells were cultivated in CGXII minimal medium containing 2 % (w/v) glucose with (“induced”) and without (“uninduced”) 100 µM IPTG using a microbioreactor cultivation system. **(B)** Growth of *C. glutamicum* ATCC 13032 in comparison to a strain lacking the *malR* gene. **(C)** For microscopic analysis, cells were grown in CGXII medium for 24 h at 30°C. Shown are wild-type *C. glutamicum* ATCC 13032 cells (row 1), *C. glutamicum* Δ*malR* cells (row 2) and *C. glutamicum* ATCC 13032 cells carrying the over expression vector pEKEx2-*malR* (row 2). The expression of *malR* is induced by the addition of 100 µM IPTG. Lipid components of the cell membrane were stained with Nile red (red); DNA was stained with Hoechst 33342 (blue). The scale bars represent 5 μm. Further microscopic pictures of cells overproducing MalR are shown in Fig. S5. **(D)** To quantify the number of cells with an altered morphology, cell counts and biovolume were analyzed using a MultiSizer 3 particle counter (Beckman Coulter, Brea, USA) equipped with a 30 µm capillary in volumentric control mode.

Fluorescence microscopy of cells stained with NileRed (lipid components) and Hoechst 33342 (DNA) revealed a heterogeneous morphology of cells overexpressing the *malR* gene. Among cells with wild type cell shape, several cells displayed a significantly elongated cell morphology upon *malR* overexpression (Fig. 3 C). The deletion mutant, however, was indistinguishable from the wild type strain (Fig. 3 C). Furthermore, overexpression of *malR* resulted in an uneven distribution of the lipid fraction as revealed by Nile red staining. The cloudy and heterogeneous distribution of the Hoechst strain also pointed towards problems regarding nucleoid condensation and segregation in the strain overexpressing *malR*. In order to quantify the observed heterogeneous morphology of cells, culture samples were further analyzed using a MultiSizer 3 particle counter (Fig. 3 D). These data show a clear shift of the cells towards an increased cell volume.

### The impact of altered MalR levels on transcription

The multitude of MalR-bound regions identified by ChAP-Seq analysis and the severe morphological changes caused by overexpression of *malR* already suggest a significant impact of MalR on the transcriptomic landscape of *C. glutamicum*. In the following, we performed a comparative transcriptome analysis of the wild type containing the empty vector pEKEx2 and the *malR* overexpressing strain using DNA microarrays. For this purpose, both strains were grown in CGXII minimal medium and harvested in the exponential phase. Additionally, we verified the obtained data using qRT-PCR with some selected samples (Fig. S4). As illustrated in Figure 4, *malR* overexpression resulted in massive changes in the global transcriptome when compared to the wild type. Overall, 170 genes showed a more than 5-fold altered mRNA level (p-value < 0.05). A complete overview on the transcriptome analysis is provided in Table S2.

**Figure 4:**
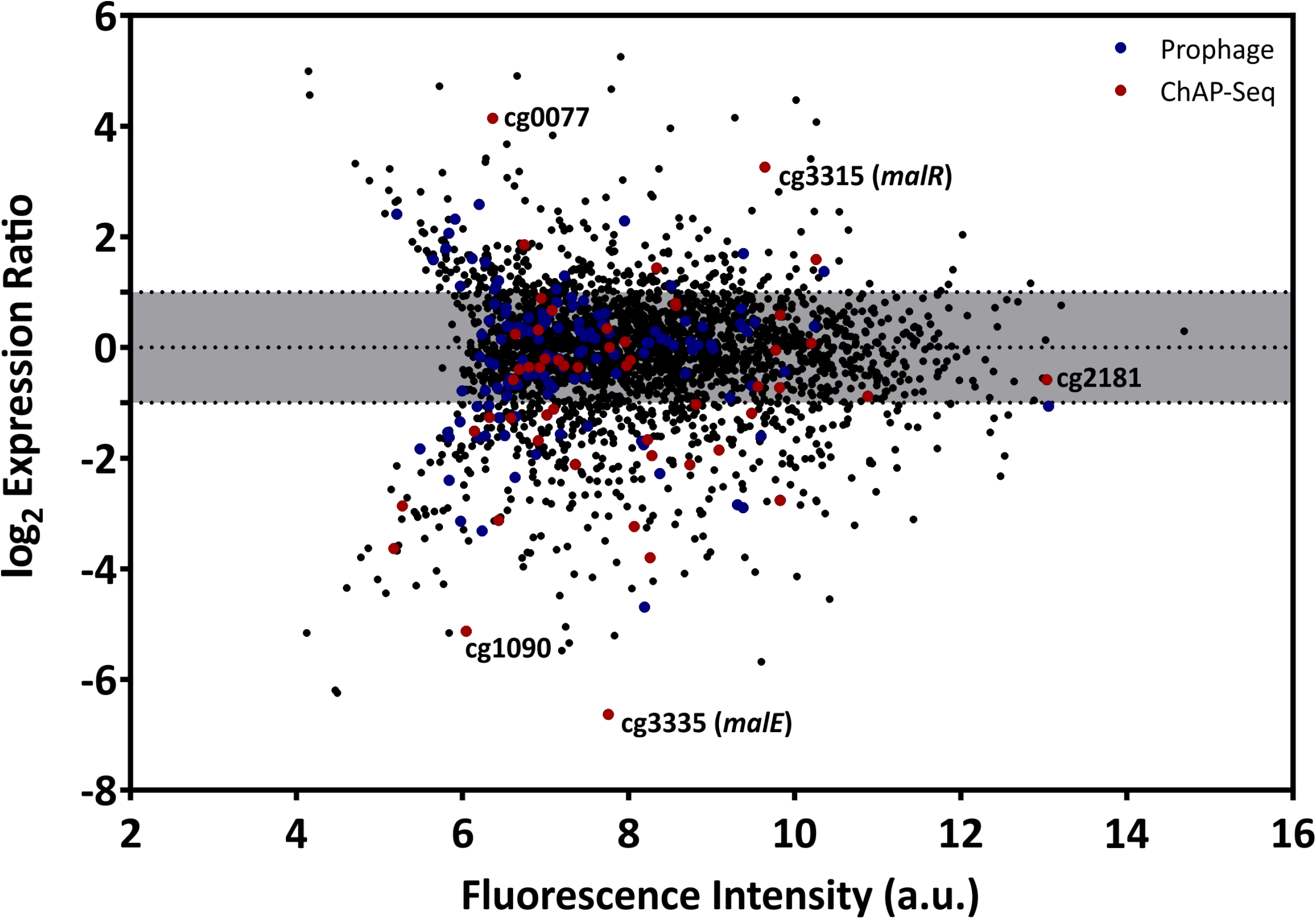
Overexpression of the *malR* gene causes global changes in the *C. glutamicum* transcriptome. Comparative transcriptome analysis of *C. glutamicum* ATCC 13032 cells carrying the overexpression vector pEKEx2-*malR* and a strain carrying the empty plasmid. Cells were cultivated in CGXII glucose minimal medium and harvested at an OD_600_ of 5. Shown is an MA-plot where the log2 of the expression ratio is plotted against the fluorescence intensity of the single spots. Red dots indicate genes that were bound by MalR in the ChAP-Seq experiment (Fig. 2).

Considering the impact of increased MalR levels on growth and cell morphology, a majority of these effects are likely the result of secondary effects. To focus on primary targets of MalR, we analyzed the impact on the expression of genes whose promoter was directly bound by MalR as found *via* ChAP-Seq analysis (selection shown in Table 1; for a complete overview see Table S1 and S2). In fact, many of the direct target genes of MalR revealed an altered mRNA level due to the overexpression of *malR*. In several cases, however, the effect was rather minor, which may be due to the chosen growth conditions lacking a specific effector molecule of the regulator. Furthermore, the majority of target genes is likely controlled by several regulatory systems impacting the transcriptional output.

**Table 1:**
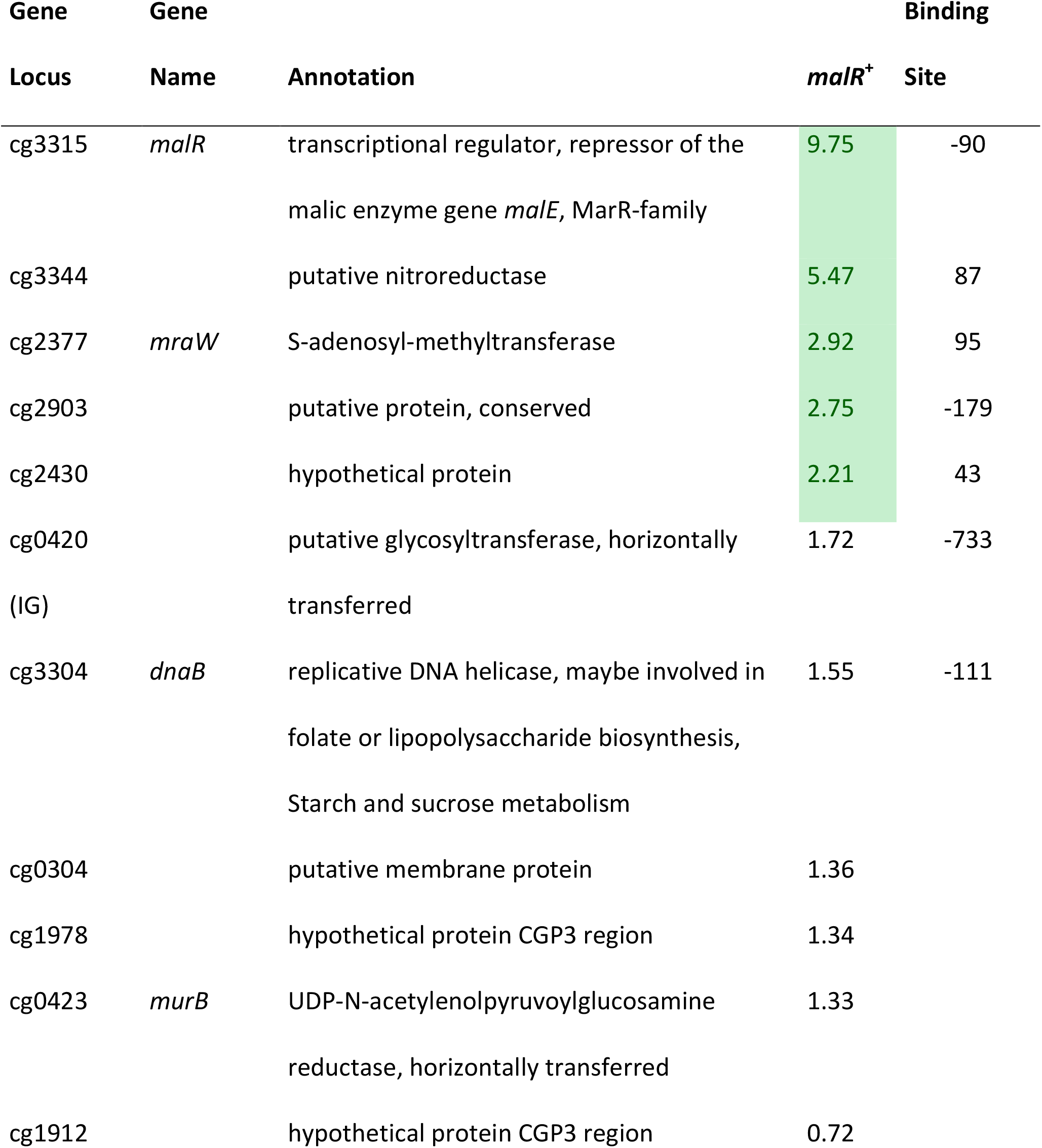
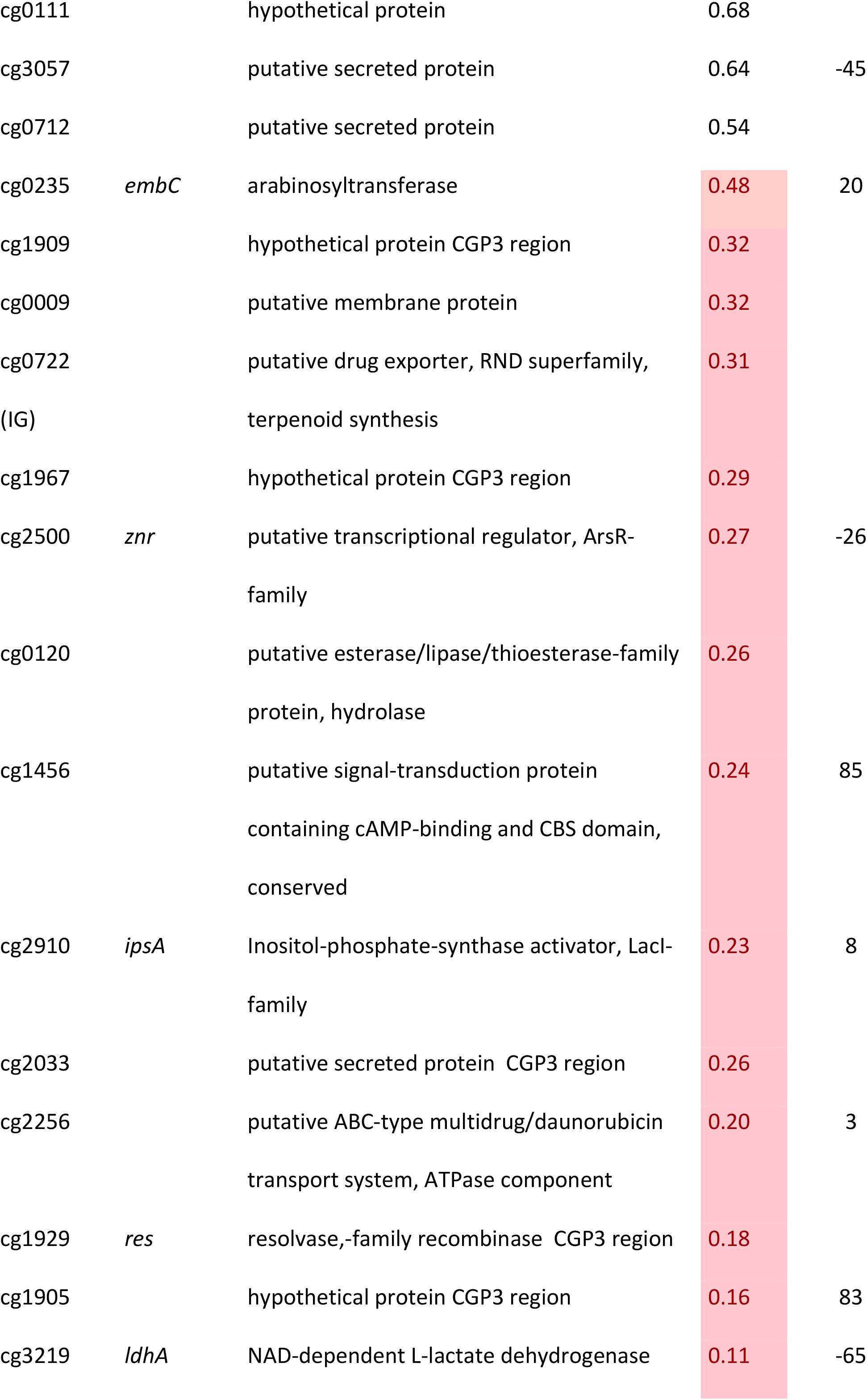
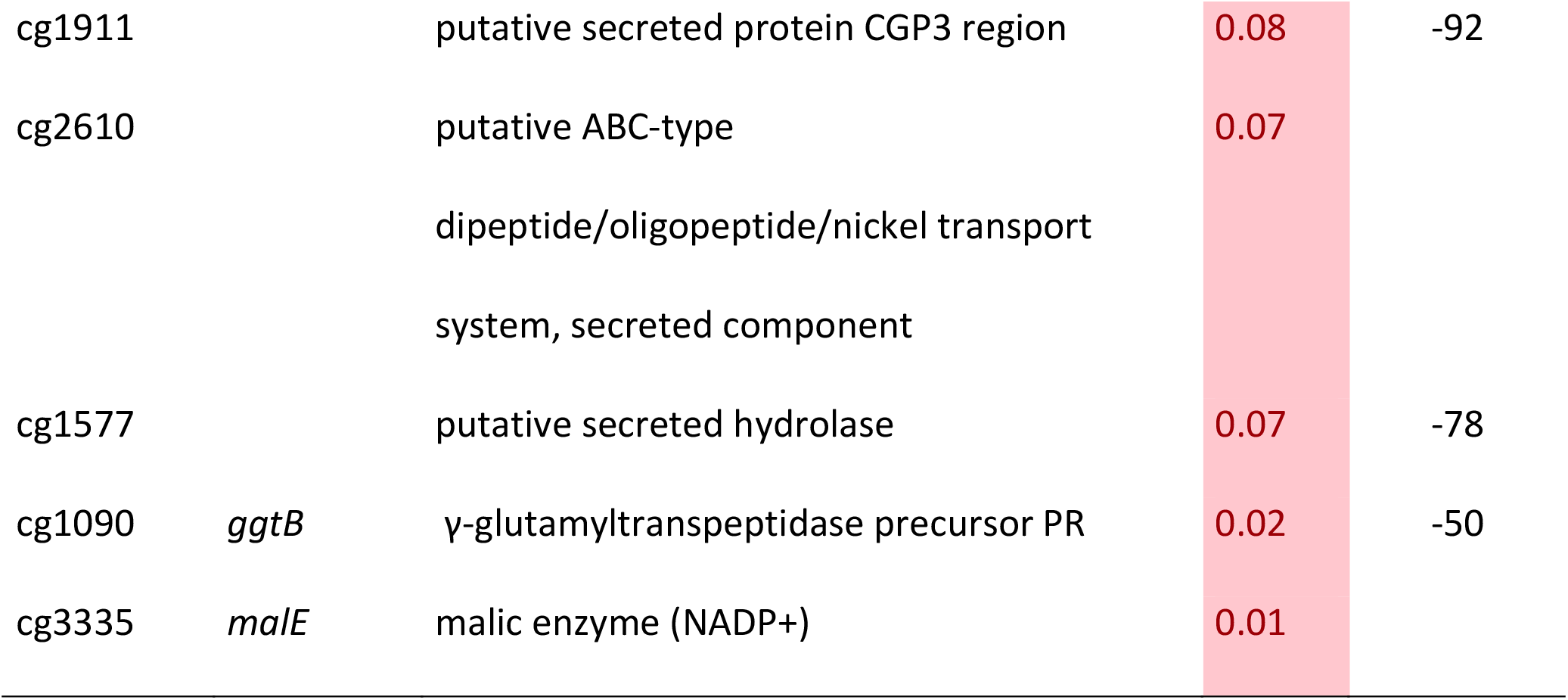
Genes bound by MalR with altered expression due to an overexpression of *malR.* Extraction of data shown in Figure 2 and Figure 4. The binding site is calculated from TSS (33) and the maximum peak position. *malR*^*+*^ indicates the fold-change of the mRNA caused by an overexpression of *malR* (p-values <0.05).

Remarkably, many genes encoding for proteins involved in cell envelope biosynthesis or remodeling were affected by *malR* overexpression. For example, the *ipsA* gene, encoding a LacI-type regulator, showed an about 5-fold downregulation in the *malR* overexpression strain. IpsA was previously described as an important regulator modulating the synthesis of inositol-derived lipids in the cell wall of *C. glutamicum* (36). Cells lacking *ipsA* revealed an elongated cell morphology with an affected growth. These findings are in line with the phenotype of the MalR overproducing strain (Fig. 3 C). Among the genes repressed by MalR are, furthermore, *embC* encoding an arabinosyltransferase involved in arabinan biosynthesis (34, 38), and several (secreted) membrane proteins of unknown function (cg0623, cg0636, cg0879, cg0952, cg1578, cg1910, cg3322). The most distinct downregulation was observed for the *malE* gene coding for the malic enzyme catalyzing the decarboxylation of malate to pyruvate. Pyruvate itself is a precursor for acetyl-CoA synthesis, a precursor molecule for fatty acid synthesis.

Among the genes showing a slightly increased expression in response to *malR* expression we found, for example, *ilvA* encoding a threonine-dehydratase that is necessary for the production of isoleucine (39). Isoleucine is a branched chain amino acid and, together with acetyl-CoA, an important precursor for the generation of branched chain fatty acids, which are part of the bacterial cell membrane. Furthermore, the *oppA* gene, coding for an oligopeptide permease required for the modulation of cell-wall associated lipids as well as mycolic acids (40), was slightly upregulated as well. Also, the expression level of methyltransferase *mraW* was about 3-fold increased. In *E. coli*, MraW was described to play an important role during cell division (41).

Among the direct targets of MalR, we also found the gene *murB*, which is involved in the synthesis of peptidoglycan building blocks by converting UDP-N-acetylglucosamine partially to UDP-N-acetylmuramic acid (42). However, its mRNA level was almost unaffected by *malR* overexpression, suggesting that further regulatory components are involved in the control of the *murAB* operon. Altogether, ChAP-Seq analysis, the impact of MalR on cell morphology and this transcriptomic study strongly emphasize an important role of MalR in the remodeling of the cell envelope.

### MalR affects the cell surface structure of *C. glutamicum*

Considering the impact of MalR on cell morphology (Fig. 3 C) we analyzed cells overproducing MalR using transmission electron microscopy (TEM) and scanning electron microscopy (SEM) (Fig. 5). Both approaches suggested differences in the cell surface structure. While wild type cells show a rather homogenous distribution in size, the strain overexpressing *malR* displayed an elongated cell morphology and significant heterogeneity with regard to cell size (Fig. 3 C and Fig. 5). Moreover, the overall cell surface structure appeared smoother. The fuzzy structure observed by TEM is also typical for the outer layer of the mycobacterial envelope (23). This electron-dense layer consists of a protein-carbohydrate matrix with only a few lipids (21).

**Figure 5:**
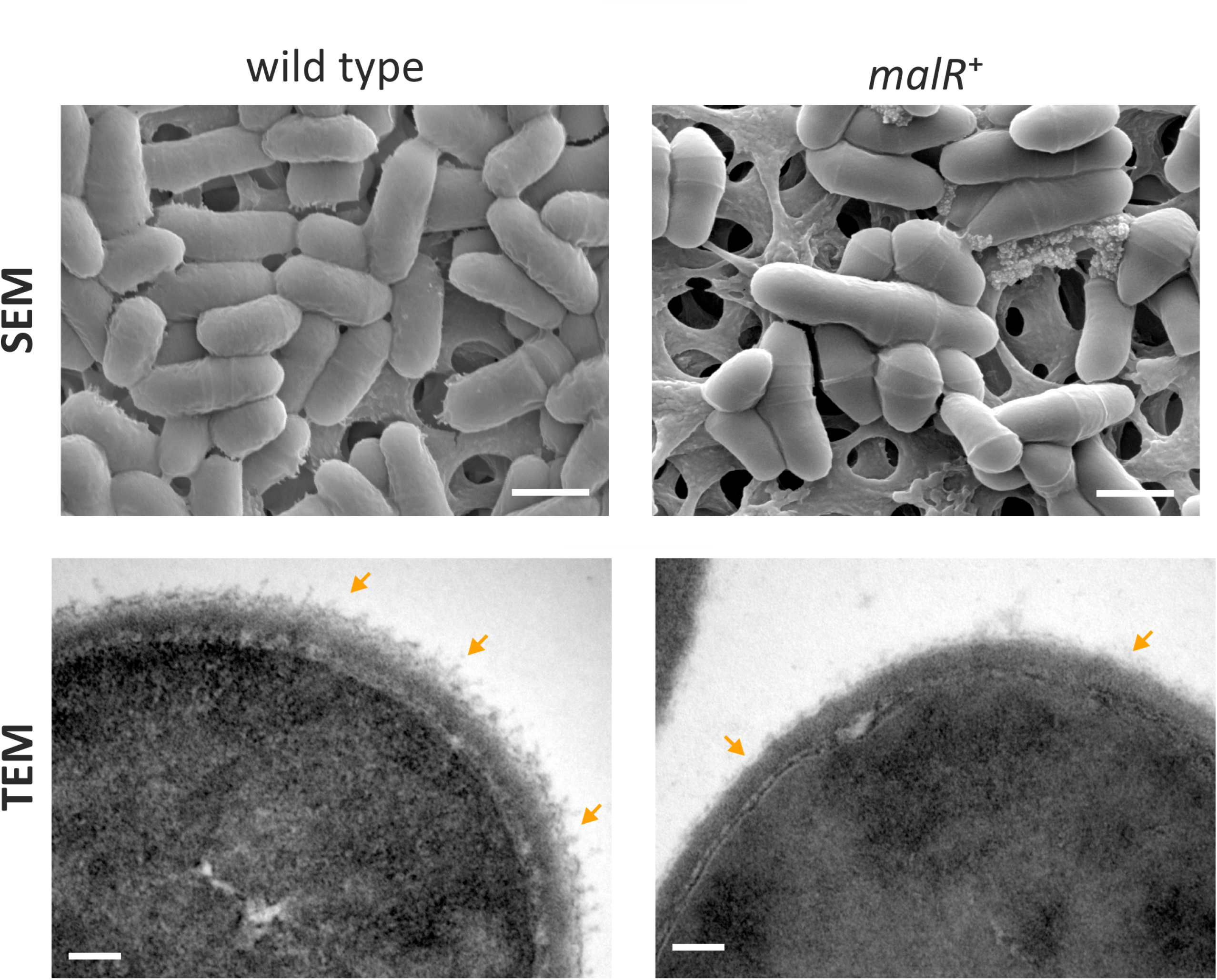
*C. glutamicum* cells overproducing MalR show an altered cell surface structure. Shown are SEM and TEM microscopy pictures of wild type cells and cells overproducing the MalR protein. SEM pictures are 15000 x magnified; TEM pictures 167000 x. For microscopic analysis, cells were cultivated in CGXII minimal medium containing 2 % (w/v) glucose and *malR* expression was induced by adding 100 µM IPTG. Cells were fixed using 3 % Glutaraldehyde in Sörensen phosphate buffer.

### MalR confers increased resistance towards β-lactam antibiotics

For a better understanding of the physiological impact of MalR, we performed phenotypic microarrays of the wild type and the Δ*malR* mutant using a Biolog system. Here, we focused on the plates PM1 and PM2A (carbon sources), PM4 (phosphorus and sulfur sources), PM9 (osmolytes), PM10 (pH), and PM11-PM13 (antibiotics). The only additives that led to a different behavior between the wild type and the *malR* deletion strain were antibiotics. To be precise, different β-lactams, tetracyclines and other examples of different substance classes revealed an altered metabolic activity of the Δ*malR* mutant (Tab. S4). Figure 6 shows four examples, emphasizing a significantly increased sensitivity of the mutant towards different cephalosporines. Whereas the wild type was able to tolerate the antibiotics at moderate levels (Amoxicillin: 3 µg / ml; Cefazolin: 0.58 µg / ml; Cephalothin: 6 µg / ml; Cefuroxime: 1.2 µg/ / ml), the Δ*malR* strain was significantly affected and did not restore metabolic activity within 40 hours under the tested conditions. An overview of all tested plates is provided in Figure S6.

**Figure 6:**
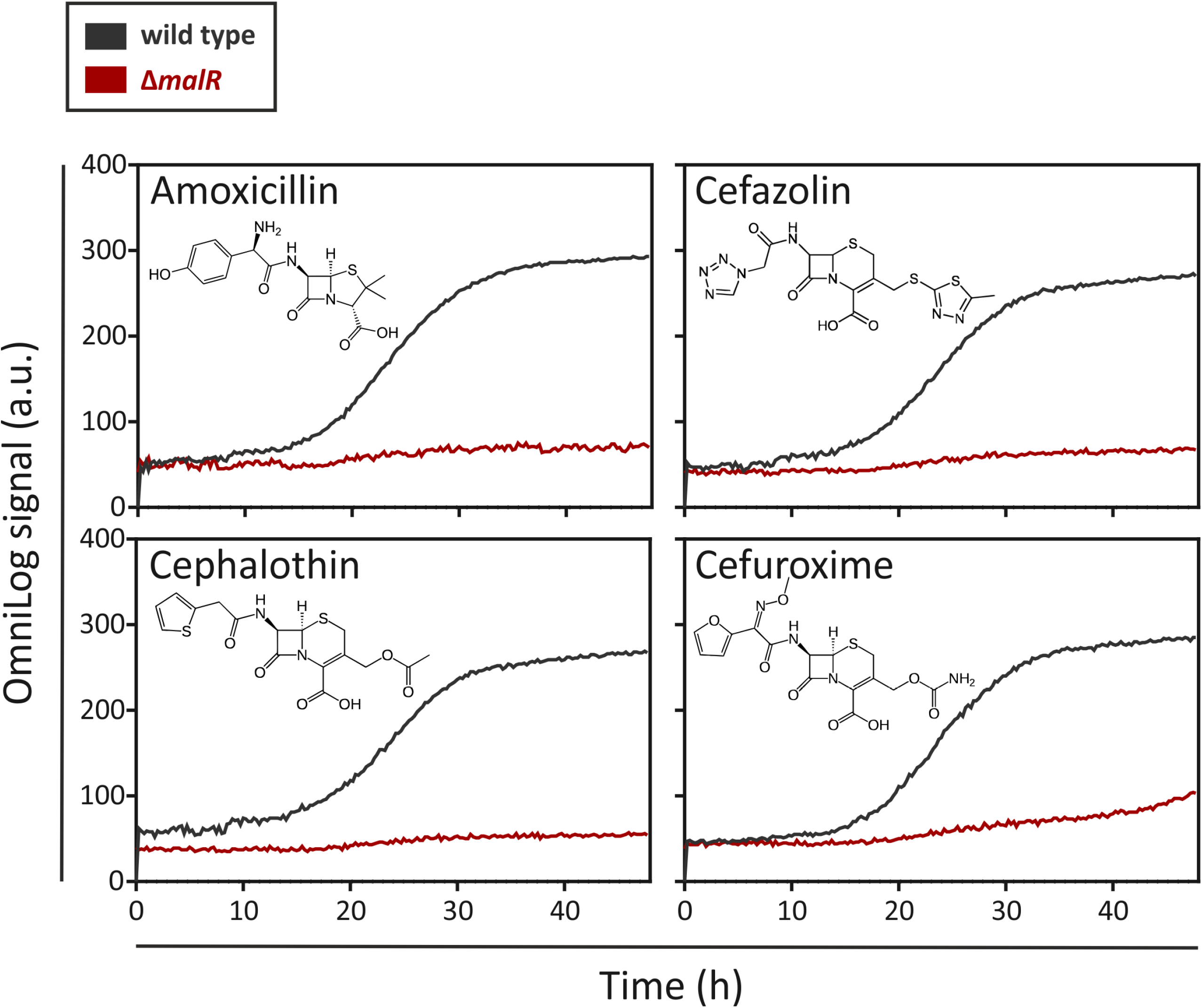
The Δ*malR* mutant strain shows a higher sensitivity towards cephalosporines in phenotypic microarrays. An OmniLog System from Biolog (Hayward CA, USA) was used to perform phenotypic microarrays with wild type *C. glutamicum* ATCC 13032 as well as the *malR* deletion strain. The experiments (PM1, PM2A, PM4, PM9, PM10, PM11, PM12, and PM13) were conducted like described in the protocol of the manufacturer (75) (42). One group of different tested conditions that led to a difference in respiration of wild type cells and Δ*malR* cells were the cephalosporins (Amoxicillin: 3 µg/ml; Cefazolin: 0.58 µg/ml; Cephalothin: 6 µg/ml; Cefuroxime: 1.2 µg/ml).

### MalR counteracts SOS-dependent induction of the CGP3 prophage

Due to several binding sites inside the CGP3 region, an impact of MalR on the inducibility of this large cryptic prophage was the focus of further experiments. For this purpose the reporter strain *C. glutamicum* ATCC 13032::P_*lys*_-*eyfp* carrying the *malR* overexpression plasmid pEKEx2-*malR* was used. The P_*lys*_-*eyfp* reporter enables the visualization of prophage induction within single cells by the production of the fluorescent protein eYFP (30). In the following, we triggered an induction of the cellular SOS response by the addition of the DNA-damaging agent Mitomycin C (MMC) and monitored its impact on CGP3 induction. The MalR level was modulated by adding increasing amounts of IPTG (10 µM, 25 µM, 50 µM). Remarkably, the fraction of CGP3 induced cells significantly declined in response to *malR* overexpression (Fig. 7 B). Also the growth of the strains was severely affected upon addiction of MMC and IPTG (Fig. 7 A). The dose responsive behavior of prophage induction in response to *malR* overexpression may suggest that MalR counteracts prophage excision under the tested conditions.

**Figure 7:**
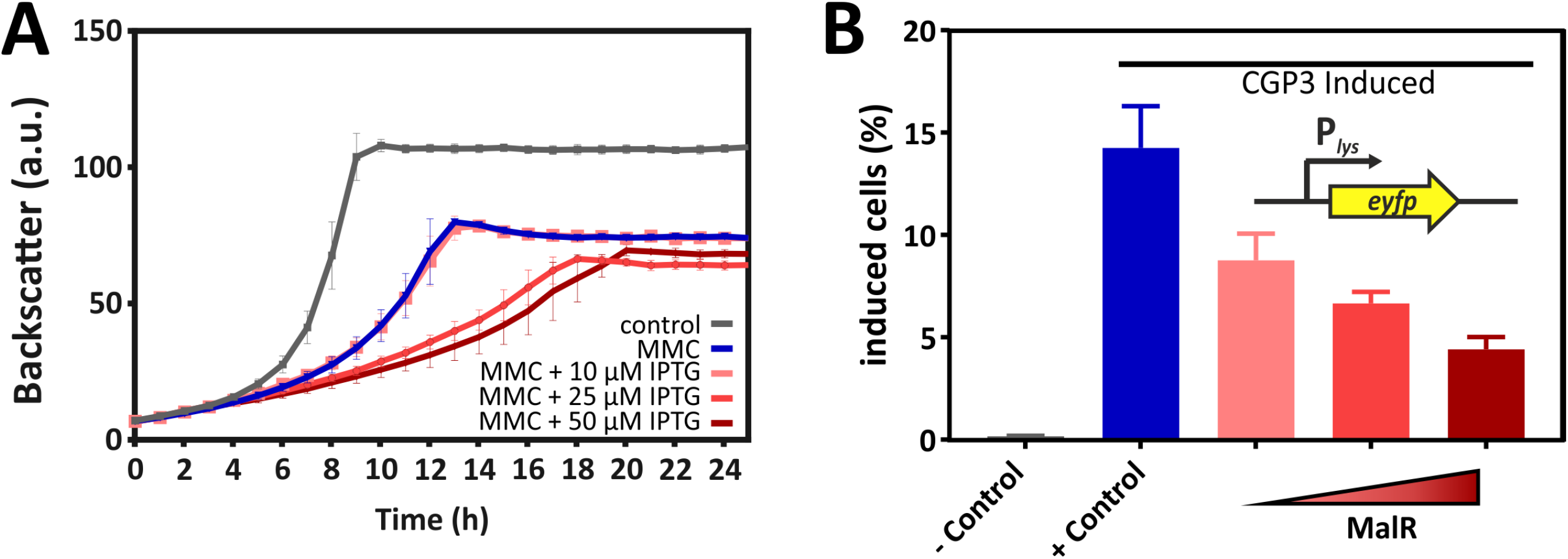
Overexpression of *malR* counteracted SOS-dependent prophage induction. *C. glutamicum* ATCC 13032::P_*lys*_-*eyfp* cells containing the plasmid pEKEx2-*malR* were cultivated in CGXII minimal medium containing 2 % (w/v) glucose (+ 25 µM Kanamycin) using a microbioreactor cultivation system. CGP3 induction was triggered by addition of 600 nM mitomycin C (MMC). For the negative control no MMC was added. Additionally, different concentrations of IPTG (0-50 µM) were used to induce expression of *malR*. Both, MMC and IPTG, were added directly at the beginning of the cultivation (0 h). Under the applied conditions, we analyzed the strain with regard to growth (A). Furthermore, the fraction of CGP3-induced cells was assessed in the stationary phase (after ∼25 hours of cultivation) using an Accuri C6 flow cytometer (B).

## 4 Discussion

With this study, we provided a comprehensive insight into the complex regulon of the MarR-type regulator MalR in *C. glutamicum*. In the last decades, members of this regulator family were rewarded with considerable attention as some MarR proteins were shown to contribute to a so-called <u>m</u>ultiple <u>a</u>ntibiotic <u>r</u>esistance phenotype (5, 7). In several cases, MarR-type regulators were described to control a small set of target genes, often located in the same operon or in divergent orientation to the regulator gene on the chromosome (7). The resulting phenotype of increased antibiotic resistance was previously proposed to be a result of decreased influx and increased efflux of the toxic compound. In this study, we now provided a genome scale profiling of MalR binding and identified more than 60 promoter regions bound by this regulator. A combination of ChAP-Seq analysis and comparative transcriptomics emphasizes MalR as a global regulator of stress-responsive remodeling of the cell envelope. Remarkably, MalR is conserved in several species of the genera *Corynebacterium* and *Mycobacterium* also including prominent pathogens like *C. diphtheriae* and *M. leprae*. Conspicuously, the genomic organization of the *malR* locus in *C. diphtheriae* is almost identical to *C. glutamicum*, where *malR* forms an operon with a gene (*uspA*) encoding a universal stress protein (26). The superfamily of Usp proteins comprises a large group of conserved proteins that can be found in all domains of life (43). In *M. bovis* BCG, the tuberculosis vaccine strain, overexpression of a particular Usp led to an increased susceptibility of the cells towards the anti-tuberculosis drug isoniazid (44). This conserved genomic organization of the *malR* locus in these species is in favor with a similar role of *C. diphtheriae* MalR in cell envelope remodeling and antibiotic resistance in this important human pathogen.

The effector molecule of MalR was so far not identified, but based on our findings, we can speculate that MalR binding is affected by one or several antibiotics and/or lipophilic compounds causing cell envelope stress.

The genome of *C. glutamicum* comprises in total nine MarR-type transcriptional regulators; four of which were already characterized in former studies. Except MalR, for none of the previously studied examples an impact on antibiotic resistance phenotype was reported so far. RosR, which constitutes a hydrogen peroxide sensitive regulator, was shown to play an important role in the oxidative stress response of *C. glutamicum* (45). The MarR-type regulator PhdR was shown to act as a repressor of the *phd* gene cluster important for phenylpropanoid utilization in *C. glutamicum*. Here, phenylpropanoids or their degradation intermediates were shown to cause dissociation of PhdR and derepression of the respective target operon (46). Finally, the isoprenoid pyrophosphate-dependent regulator CrtR was recently described as being involved in the regulation of carotenoid biosynthesis and thus represents an example for a rather specialized MarR-type regulator (47). MalR itself was firstly reported by Krause *et al*. as a repressor of the *malE* gene encoding an NADP^+^-dependent malic enzyme in *C. glutamicum* (31). This role is supported by our study, where *malE* was among the genes most affected by MalR overexpression. MalE catalyzes the decarboxylation of malate to pyruvate while generating NADPH (48). Pyruvate, a precursor for acetyl-CoA synthesis, as well as NADPH as reducing agent, are required for fatty acid biosynthesis (49). For different oleaginous microorganisms it is known, that malic enzymes play a crucial role in lipid generation (50–52). For example, two malic enzymes were recently shown to be important for triacylglycerol and antibiotic production in *Streptomyces coelicolor* (53).

Genome-wide profiling of MalR-bound DNA using ChAP-Seq analysis, unraveled more than 60 direct target genes and operons in addition to *malE*. Thus, the global impact of MalR ranges from peptidoglycan biosynthesis (*murA*-*murB* (42)), to the synthesis of arabinogalactan (*embC* (34)), and cell wall associated lipids and mycolic acids (e.g. via *oppA* and *ipsA*). For example, the *oppA* gene codes for an oligopeptide permease, which was further characterized in *M. tuberculosis* (40). Flores-Valdez and others could show that OppA is required for the modulation of cell-wall associated lipids as well as mycolic acids. The LacI-type regulator IpsA, which is itself repressed by MalR, was previously shown to trigger the expression of the *ino1* gene encoding the inositol phosphate synthase. Deletion of *ipsA* resulted in a severe decrease of inositol-derived lipids and an abolished mycothiol biosynthesis (36). Remarkably, a Δ*ipsA* mutant features a similar cell morphology as a *malR* overexpression suggesting that some phenotypic effects may be indirectly resulting from reduced IpsA levels (Fig. 3 C). An overview on the function of selected MalR targets is provided in the model shown in Figure 8.

**Figure 8:**
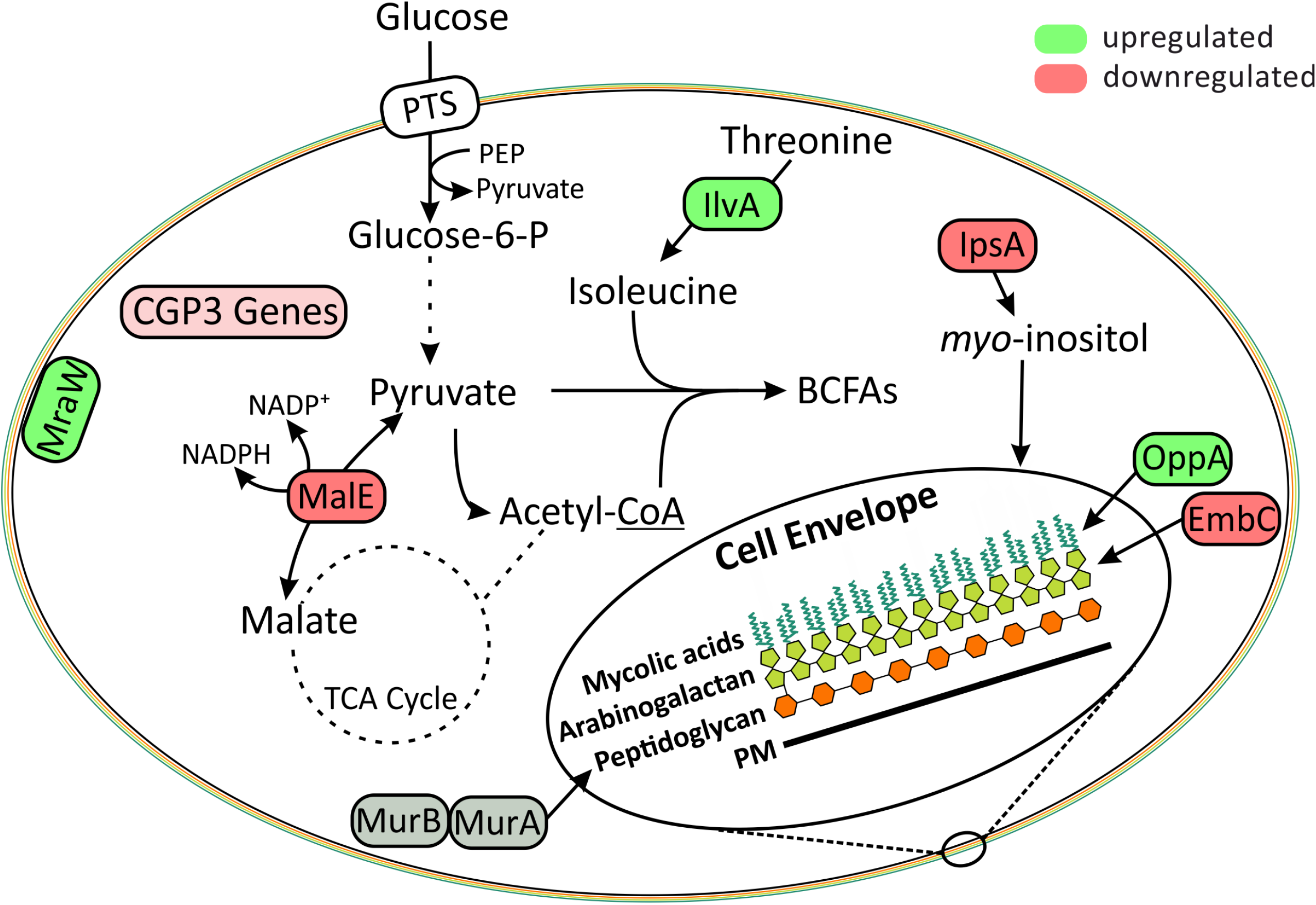
Model of the regulatory circuit influenced by the overproduction of MalR. The proteins shown are transcriptionally either up (green) or down regulated (red) in a *C. glutamicum* strain overproducing MalR. MurA and MurB are not significantly regulated under the chosen conditions but also represent a direct target of MalR (Fig. 2). Additionally, the majority of CGP3 genes were downregulated (light red) in our study. The displayed model emphasizes an important role of MalR in stress-responsive cell envelope remodelling of *C. glutamicum*.

A role in cell envelope remodeling appears to be a common theme in the family of MarR regulators. For example, the MarR-type regulator SlyA was shown control several targets impacting cell envelope composition, some of which have direct implications on the resistance of *Salmonella enterica* towards antibiotics (e.g. polymyxins) or virulence (54). The regulator Rv1404 from *Mycobacterium tuberculosis* was also shown to contribute to an adaptation of the cells to the host environment by enhancing the acid-tolerance of this bacterium (55). Altogether, these examples emphasize different roles of MarR-type regulators. Whereas some proteins appear to conduct very distinct regulatory functions, like the control of a certain catabolic gene cluster, others (like MalR or SlyA) act as global regulators orchestrating complex adaptive strategies in response to environmental stresses.

A further prominent target of MalR appeared to be the cryptic prophage element CGP3. MalR bound to overall 13 regions within the CGP3 element and overexpression of the *malR* gene counteracted CGP3 induction upon addition of the SOS inducing antibiotic mitomycin C. So far, little is known about the role of MarR-type regulators in the control of horizontally acquired elements. In *Bacillus subtilis*, a *pamR* deficient strain displayed altered expression of prophage genes, however the precise regulatory impact remained unclear (56). Another example is depicted by RovA, which is a MarR-type transcription factor in *E. coli*, *Yersinia pseudotuberculosis* and its homologue SlyA from *Salmonella* (57). Former studies revealed that RovA and SlyA act as countersilencers of H-NS target promoters controlling genes impacting virulence in *Yersinia and Salmonella species* (54, 58–60). We previously reported on the xenogeneic silencer CgpS, which plays a crucial role in the silencing of the CGP3 island by binding to AT-rich DNA (61). MalR itself binds to an AT-rich palindromic motif whose composition, of course, increases the likelihood of an overrepresentation in horizontally-acquired regions. However, the precise regulatory impact of MalR on CGP3 activation remains unclear. The observed reduction of CGP3 induction in response to *malR* overexpression speaks against a countersilencing mechanism as reported for RovA. In contrast, the majority of phage targets appeared to be repressed by MalR. In physiological terms, a repressive function of MalR towards CGP3 genes could be overcome by the presence of an effector molecule leading to a dissociation of a putative MalR dimer. So, we could speculate on a role of MalR in stress-responsive induction of the CGP3 prophage, which could literally leave the “sinking ship” when the life of its host is threatened by harsh environmental conditions.

With this study, we provide a comprehensive overview on the many targets controlled by MalR and suggest an important function of this MarR-type regulator in the coordinated control of genes with an impact on cell envelope composition. The relevance of this global response is reflected by the severely increased sensitivity of a *malR* mutant to several β-lactam antibiotics and is further supported by several other cases where members of this family contributed to enhanced antibiotic resistance (7, 11, 13, 54). We have gained a first glimpse on a complex adaptive response. However, many - if not most - of the MalR targets encode proteins of unknown function. For several others only very limited data are available. So, many more studies are needed to understand the molecular principles behind this adaptive response. With the multitude of targets identified in this study, we provide a starting point for further studies aiming to enhance our understanding of bacterial adaptation to stress.

## 5 Material and Methods

### Bacterial strains, plasmids and growth conditions

All bacterial strains and plasmids used in this work are listed in Table 2. For cloning and plasmid construction the strain *E. coli* DH5α was used, whereas the strain *E. coli* BL21 was used for protein production. These strains were – unless stated otherwise – cultivated in lysogeny broth (LB, (62)) containing 50 µg/ml Kanamycin. *Corynebacterium glutamicum* ATCC 13032 was used as a wild type strain (26). All derived *C. glutamicum* strains were cultivated in brain heart infusion medium (BHI, Difco Laboratories, Detroit, MI, USA) or in CGXII minimal medium containing 2% (w/v) glucose (63); if necessary 25 µg/ml Kanamycin was added.

**Table 2:**
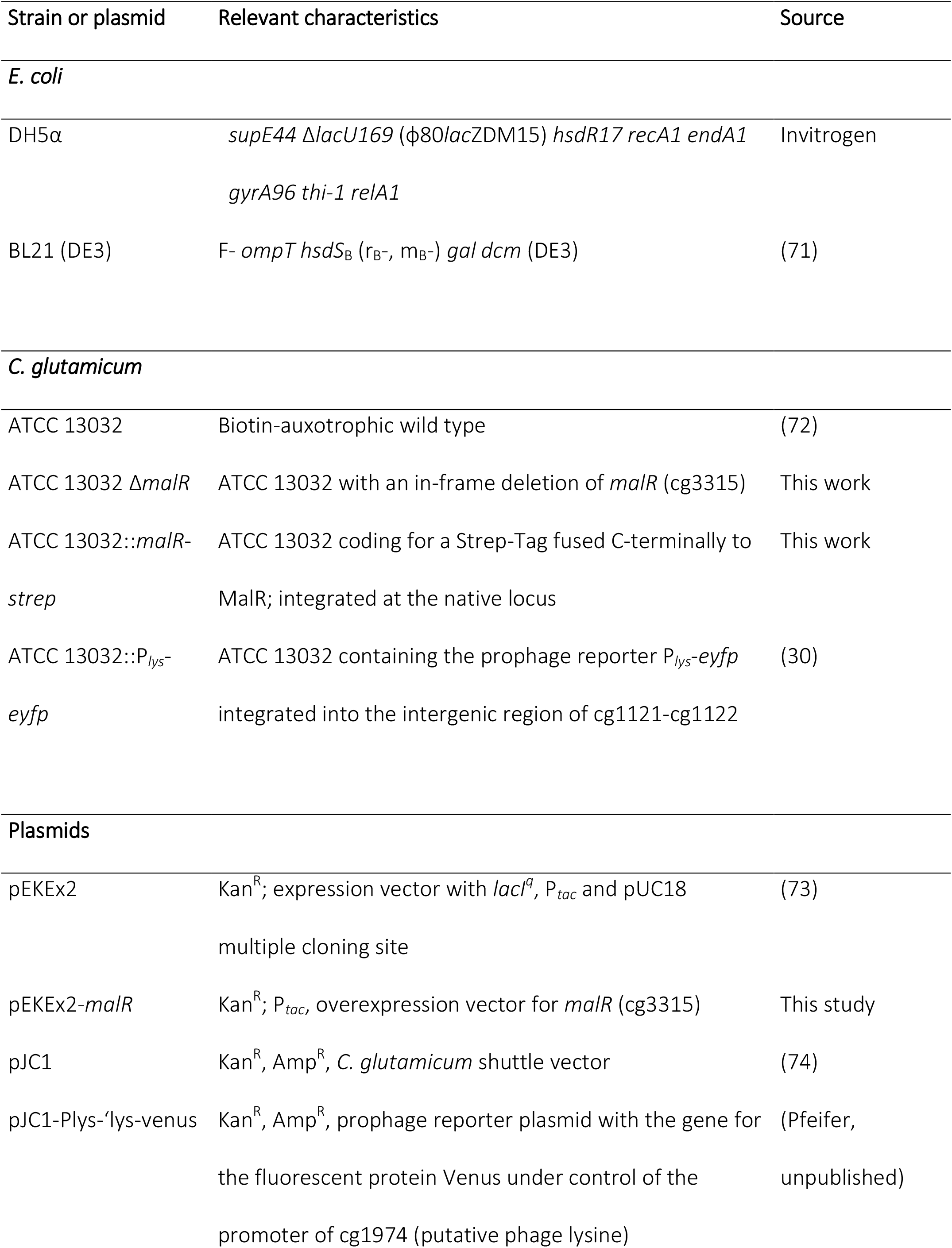
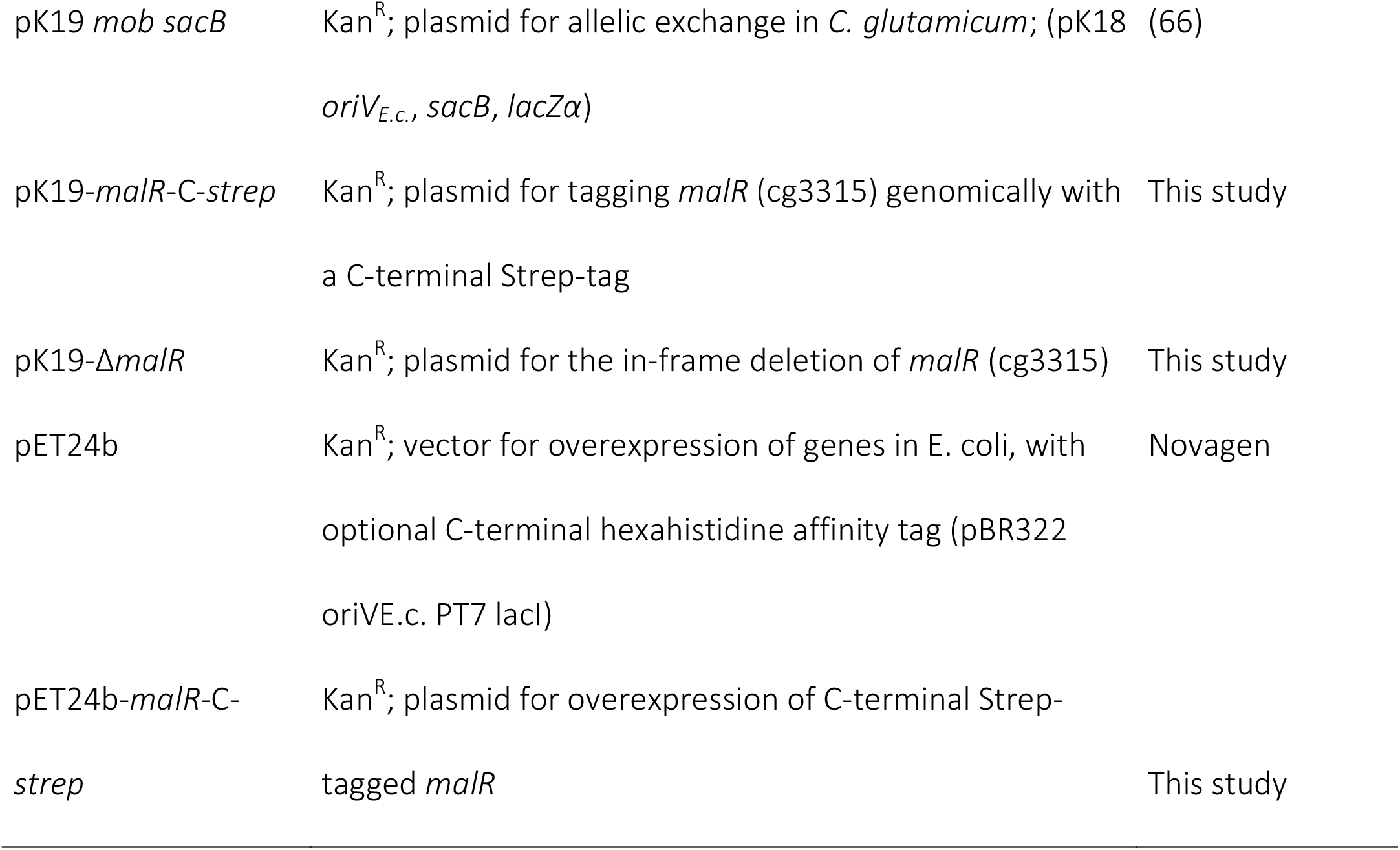
Strains and plasmids used in this work.

Growth experiments were conducted in the BioLector microbioreactor system (m2p labs, Baesweiler, Germany) (64). Therefore, 750 µl CGXII medium containing 2% (w/v) glucose and the particular stated additives (e.g. Isopropyl β–D-1-thiogalactopyranoside (IPTG)) were inoculated to an OD_600_ of 1 in 48-well microtiter plates (Flowerplates, m2p labs) and cultivated for at least 24 h at 30°C and 1200 rpm shaking frequency. Fluorescence as well as backscatter measurements were taken every 15 minutes.

### Recombinant DNA work and construction of chromosomal insertions or deletions

All standard laboratory methods (PCR, DNA restriction, Gibson Assembly) were performed according to standard protocols and manufacturer’s instructions (62, 65). The used oligonucleotides as well as details regarding the plasmid construction are provided in the supplementary information (Tab. S3 A and S3 B).

For chromosomal integration or deletion, a two-step homologous recombination system based on the suicide vector pk19*mobsacB* was used (66, 67). This vector contained 500 bps of each site flanking the targeted sequence in the genome of *C. glutamicum*.

### Chromatin Affinity Purification and next generation sequencing (ChAP-Seq)

Pre-cultures for ChAP-Seq were conducted like described above. As a main-culture, *C. glutamicum* ATCC 13032::*malR-strep* was grown in CGXII medium containing 2% (w/v) glucose for 5 hours at 30°C and 120 rpm shaking frequency. The subsequent preparation of ChAP-Seq samples and the implementation of this method was conducted as described by Pfeifer et al. (61).

### Protein purification

MalR with a C-terminal Strep-tag was heterologously produced in *E. coli* BL21 (DE3), transformed with pET24b-*malR*-C-strep. Cells were grown to an OD_600_ of 0.6 at 37°C as described above. Subsequently, the protein production was induced using 100 µM IPTG and the cultivation was continued for 5 h at 16°C. Cells were harvested by centrifugation for 15 min at 5300 x *g* and 4°C and the pellets were snap-frozen using liquid nitrogen. For cell disruption the pellets were thawed on ice and resuspended in buffer A (50 mM Tris-HCl, 1mM EDTA, pH 8.0) containing *cOmplete* Protease Inhibitor (Roche, Basel, Switzerland). This cell suspension was then treated with a French pressure cell for three passages at 172MPa. Cell debris was removed by centrifugation for 30 min at 5300 x *g* and 4°C and a subsequent ultracentrifugation for 1h at 150,000 x *g* and 4°C. The tagged protein was then purified with a 1-ml bed volume Strep-Tactin-Sepharose column (IBA, Göttingen, Germany), following the manufacturer’s protocol.

### Electrophoretic mobility shift assays

The investigation of the binding properties of MalR and as an *in vitro* verification of the results obtained by ChAP-seq analysis, electrophoretic mobility shift assays (EMSAs) were performed. Therefore 100 bp DNA fragments centering the peak maximum of each particular promoter region were amplified using PCR (oligonucleotide sequences are given in the supplementary informations, Tab. S3 C) and analyzed and purified using an agarose gel with subsequent gel extraction with the “PCR clean-up and Gel extraction” Kit from Macherey-Nagel (Düren, Germany). A total of 90 ng of DNA per lane was incubated with different molar excesses of purified MalR protein (3-fold and 10-fold molar excess) for 30 min in bandshift-buffer (50 mM Tris-HCl, 5 mM MgCl_2_, 40 mM KCl, 5 % (v/v) glycerol, pH 7.5). Subsequently, samples were separated using a 10% native polyacrylamide gel electrophoresis as described previously (61).

### Transcriptome analysis using DNA Microarrays

To analyze the global transcriptomic alterations triggered by an overexpression of *malR C. glutamicum* ATCC 13032 cells harboring either the empty vector pEKEx2 or the overexpression plasmid pEKEx2-*malR* were cultivated in CGXII medium containing 2% (w/v) glucose and 100 µM IPTG as described previously. Subsequently, total RNA of these cultures was prepared using the RNeasy Mini Kit (QIAGEN, Hilden, Germany) following the manufacturers protocol. The cDNA labeling, and the DNA microarray analysis was performed as described previously (68). The array data have been deposited in the GEO database (http://ncbi.nlm.nih.gov/geo) under the accession number: GSE116655.

### Verification of the transcriptomic data measuring mRNA levels by quantitative real-time PCR (qRT-PCR)

Preparation of total RNA from *C. glutamicum* cultures was carried out as described above. Measurement of differential gene expression was conducted using a qTower 2.2 (Analytik Jena, Jena, Germany) with the Luna^®^ Universal One-Step RT-qPCR Kit (New England Biolabs, Ipswitch, USA). Primer pairs used for the analysis are listed in Table S3 D. For all samples 100 ng of total RNA were used as a template and all measurements were performed in biological as well as in technical duplicates. The Ct-values of the samples were obtained using qPCR-soft 3.1 (Analytik Jena). Subsequently, the relative transcriptional changes were calculated using the following equation:

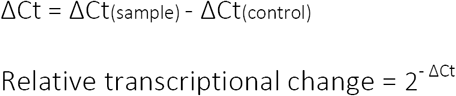

### Fluorescence microscopy and staining

For microscopic analysis, cells were cultivated in CGXII medium containing 2% (w/v) glucose (as described above) and grown for 24 h at 30°C. Lipids were stained with Nile red and DNA was stained with Hoechst 33342 (Sigma-Aldrich, Munich, Germany). Therefore 10 µl of the cell suspensions were centrifuged for 5 min at 8,000 x *g* and the pellet was resuspended in 500 µl PBS containing 100 ng/ml Hoechst 33342 and 250 ng/ml Nile red. The cells were incubated for 30 min at room temperature and subsequently analyzed microscopically using an AxioImager M2 (Zeiss, Oberkochen, Germany) with a Zeiss AxioCam MRm camera and a Plan-Apochromat 100x, 1.40 Oil phase contrast oil-immersion objective. Fluorescence was measured using the 63 HE filter for Nile red fluorescence and the filter set 49 for Hoechst fluorescence.

The optimal exposure time for the different fluorescence images was determined with the automatic measurement option of the AxioVision Rel. 4.8 software (Carl Zeiss MicroImaging GmbH) and the pictures were analyzed with the same software.

### Scanning and transmission electron microscopy

For scanning (SEM) and transmission electron microscopy (TEM) cells were treated as described previously (69). Further details on the cultivation preparation procedure are given in the supplementary information.

### Cell counting and determination of the cell size

Cell counts and biovolume were analyzed using a MultiSizer 3 particle counter (70) equipped with a 30 μm capillary in volumetric control mode. For the measurement, cells were grown for 24 hours as described above and afterwards diluted to an OD_600_ ≤ 0.025 in CASYton buffer (Schärfe Systems, Reutlingen, Germany). Only particles sizing from 0.633 µm to 18 µm were analyzed.

### Phenotypic analysis of *C. glutamicum* Δ*malR* and *C. glutamicum* ATCC 13032

Phenotypical characterization of the strains *C. glutamicum* Δ*malR* and *C. glutamicum* ATCC 13032 was performed using the Phenotypic MicroArrays from BIOLOG (Biolog Inc., Hayward, CA, USA). Both strains were compared regarding their respiratory activity in the presence of various carbon sources (PM1, PM2), phosphorus and sulfur sources (PM4), different osmolytes (PM9), pH-values (PM10) and antibiotics (PM11, PM12, PM13). Experimental setup was carried out as described in the BIOLOG protocol for analysis of *Bacillus subtilis* and other Gram-positive bacteria (Biolog Inc., Hayward, CA, USA). In short, both strains were grown overnight at 30°C on blood agar plates. Cells were inoculated in the different PM-media containing 1% (v/v) of the redox dye (dye mix F) to a turbidity of 81% transmittance. Afterwards each well of the PM-plates was filled with 100 µl of the corresponding inoculation medium. Phenotypic MicroArrays were analyzed using the OmniLog incubator (Biolog Inc., Hayward, CA, USA) at 30°C for 48 hours with a measuring rate of 15 minutes. Data visualization was performed by the BIOLOG software OM_Pl_Par 1.20.02 for parametric analyses. For selected examples GraphPad Prism 7 was used for visualization. An overview of all results is shown in Figure S6.

### Flow cytometric analysis

The CGP3 prophage induction was assessed by flow cytometric analysis of a *C. glutamicum* strain harboring a genomically integrated prophage reporter (ATCC 13032::P_*lys*_-*eyfp*) using a BD Accuri C6 flow cytometer (BD biosciences, Heidelberg, Germany). Cells were cultivated in CGXII medium containing 2% (w/v) glucose (as described above) and grown for 25 h at 30°C. As appropriate, different concentrations of IPTG (10 µM, 25 µM and 50 µM) as well as 600 nM Mitomycin C were used. Flow cytometric analysis was performed using a 488 nm laser and a 530/30 nm filter for measuring eYFP fluorescence. In total, 100,000 events were analyzed per sample and data was analyzed using BD Accuri C6 software and visualized using GraphPad Prism 7. The gating was performed according to the uninduced negative control.

## 6 Acknowledgements

We thank Tino Polen and Doris Rittmann for assistance in sequencing and comparative transcriptome analysis, Holger Morschett for help with the Coulter Counter, Cornelia Gätgens for technical support, and Eugen Pfeifer and Marc Keppel for fruitful discussions.

## 10 Supporting Information

Methods Supplementary information on methods used in this study

Table S1: DNA regions discovered as binding sites from MalR via ChAP-Seq

Table S2: Transcriptomic changes triggered by an overproduction of MalR

Table S3 A: Oligonucleotides used in this study

Table S3 B: Construction of plasmids used in this study

Table S3 C: Fragments for EMSAs

Table S3 D: Oligonucleotides for qRT-PCR analysis

Table S4: Antibiotics affecting the growth of wild type cells or Δ*malR* cells during phenotype microarrays (BioLog)

Figure S1: Prediction of the secondary structure of MalR

Figure S2: *In vitro* DNA-binding of purified MalR protein Figure S3: Extracted binding motifs of MalR

Figure S4: Verification of DNA-microarray data showing transcriptomic changes due to overproduction of MalR using qRT-PCR

Figure S5: MalR overproduction causes severe growth defects of *C. glutamicum*

Figure S6: Overview of all tested PM plates for a comparison of *C. glutamicum* Δ*malR* and *C. glutamicum* wild type

