## Supplementary material for "The MarR-type regulator MalR is involved in stress-responsive cell envelope remodeling in *Corynebacterium glutamicum*": Figure S1

### Supporting Information

#### Figure S1

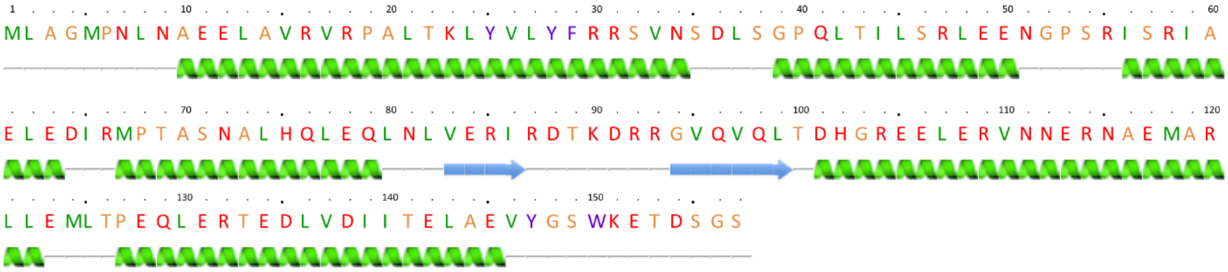


**Figure S1: Prediction of the secondary structure of MalR.** Using the online tool Phyre[2], the secondary structure of MalR was predicted with 99.9% confidence of 92 % of residues (Kelly L.A., Mezulis S., Yates C., Wass M., Sternberg, M. 2015. Nat. Protoc.10:845–858.).
