## Supplementary material for "The MarR-type regulator MalR is involved in stress-responsive cell envelope remodeling in *Corynebacterium glutamicum*": Figure S2

### Supporting Information

#### Figure S2

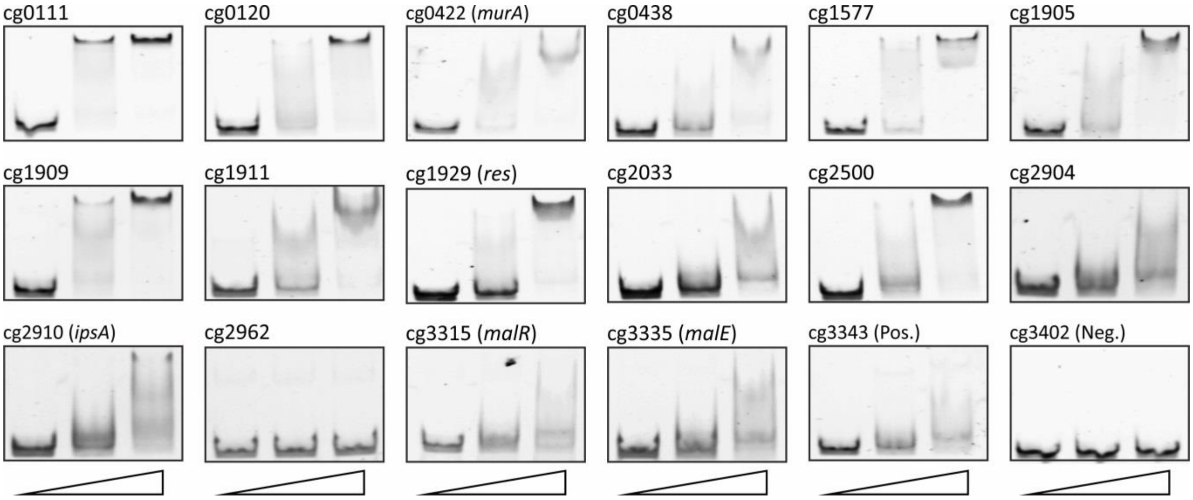


**Figure S2:** ***In vitro*** **DNA-binding of purified MalR protein.** Electrophoretic mobility shift assays (EMSAs) were performed to verify binding of MalR to promoter regions identified via ChAP-Seq. Therefore, MalR was purified with a C-terminal Strep-tag fusion. At total, 90 ng of 100 bp DNA fragments (50 bp up- and downstream the peak maximum) were incubated without protein (first lane), with 3 molar excess (228 nM, second lane) and 10 molar excess of purified MalR (760 nM, third lane) for 30 min in bandshift-buffer (50 mM Tris-HCl, 5 mM MgCl2, 40 mM KCl, 5 % (v/v) glycerol, pH 7.5). Subsequently, samples were separated on a 10% native polyacrylamide gel electrophoresis and gels were stained using SYBR Green I Nucleic Acid Gel Stain (Lonza, Rockland, ME, USA).
