## Supplementary material for "The MarR-type regulator MalR is involved in stress-responsive cell envelope remodeling in *Corynebacterium glutamicum*": Figure S3

### Supporting Information

#### Figure S3

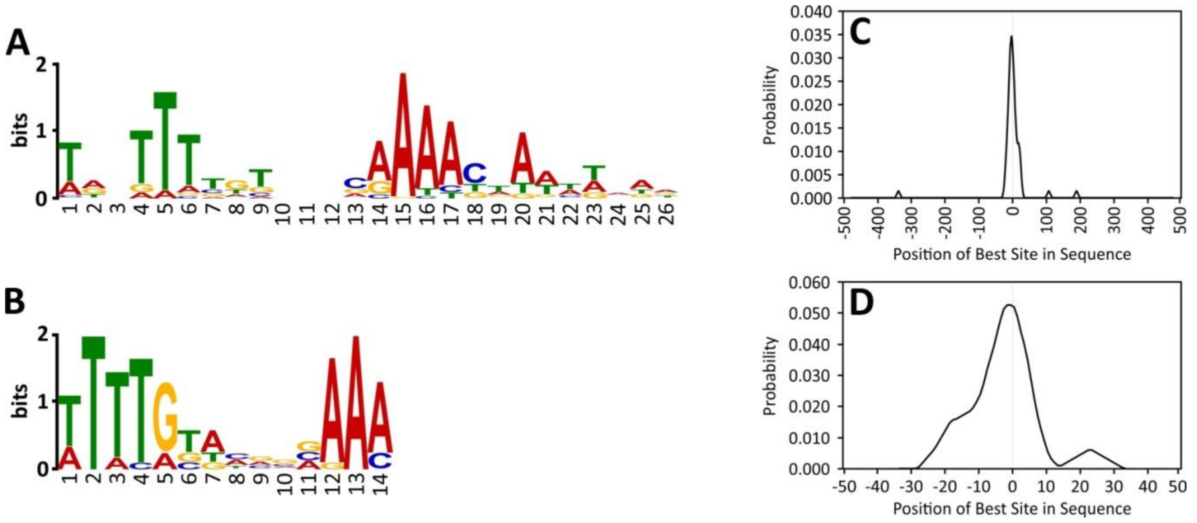


**Figure S3: Extracted binding motifs of MalR** The online tool MEME-ChIP (Ma W, Noble WS, Bailey TL. 2014. Nat Protoc 9:1428–1450.) was used to predict MalR binding sites on the basis of all targets sequences (each sequence 1000 bp) identified via ChAP-Seq analysis **(A)** and based on the 16 MalR targets (100 bp each) verified using EMSAs **(B)**. **(C, D)** The distribution of the predicted binding sites throughout the single uploaded DNA sequences was determined using CentriMo (Bailey TL, MacHanick P. 2012. Nucleic Acids Res 40:e128.).
