## Supplementary material for "The MarR-type regulator MalR is involved in stress-responsive cell envelope remodeling in *Corynebacterium glutamicum*": Figure S4

<u>Supporting Information</u>

<u>Figure S4</u>

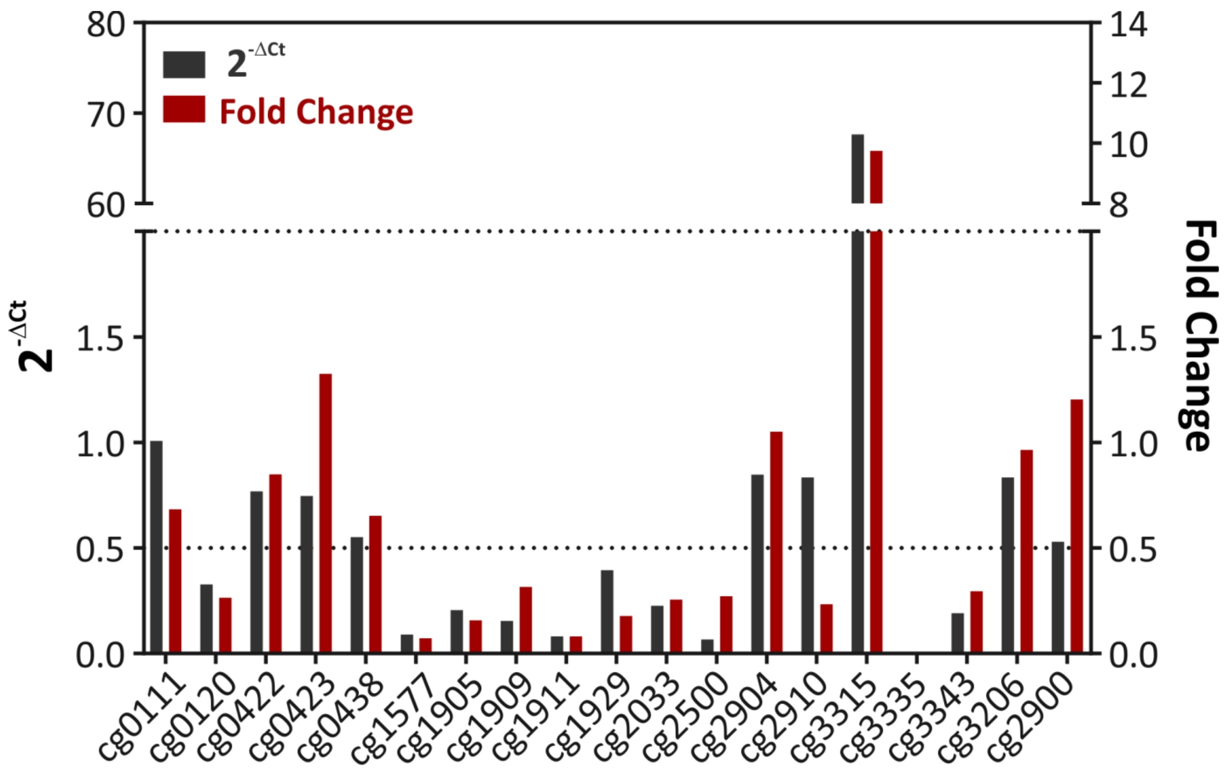


**Figure S4: Verification of DNA-microarray data using qRT-PCR.** Microarray data were verified with qRT-PCRs (oligonucleotides are listed in Table S3 D). The dark-grey bars represent the relative transcriptional change based on the qRT-PCR data and the red bars show the average fold-change obtained from DNA microarray experiments. The values below 0.5 are classified as "downregulated", the values above 2 are classified as "upregulated". In case of cg3335 no bars can be detected because of very low values (qRT-PCR: 0.018, DNA-microarray: 0.007).
