## Supplementary material for "The MarR-type regulator MalR is involved in stress-responsive cell envelope remodeling in *Corynebacterium glutamicum*": Figure S5

### Supporting Information

#### Figure S5

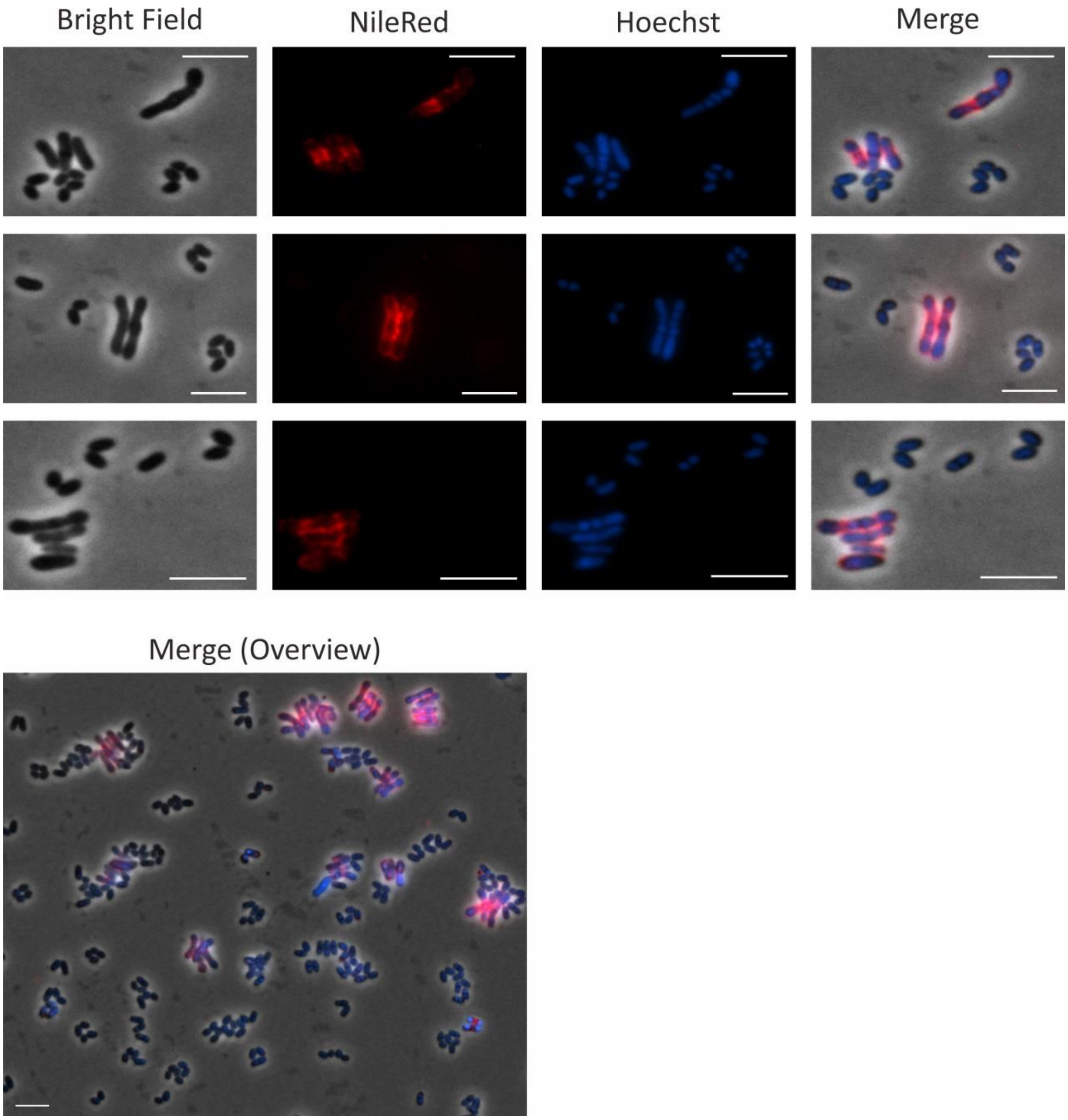


**Figure S5: MalR overproduction causes severe growth defects of *C. glutamicum*.** For microscopic analysis, cells were grown in CGXII medium for 24 h at 30°C. Shown are *C. glutamicum* ATCC 13032 cells carrying the over expression vector pEKEx2-*malR*. The expression of *malR* is induced by the addition of 100 µM IPTG. Lipid components of the cell membrane were stained with Nile Red (red); DNA was stained with Hoechst 33342 (blue). The white scale bars represent 5 µm. In addition to chosen examples of elongated cells an overview is shown to present the distribution of elongated cells inside the samples.
