## Supplementary material for "The MarR-type regulator MalR is involved in stress-responsive cell envelope remodeling in *Corynebacterium glutamicum*": Figure S6

### Supporting Information

#### Figure S6

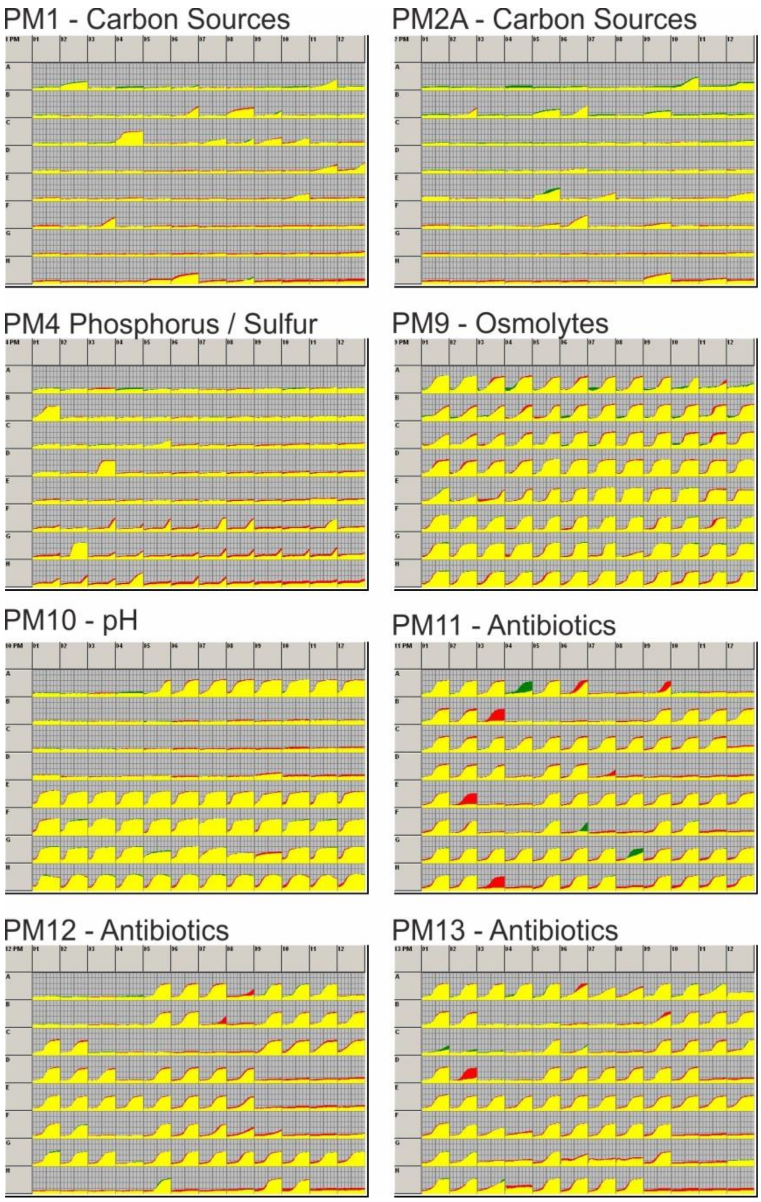


**Figure S6: Overview of all tested PM plates for a comparison of *C. glutamicum* Δ*malR* and *C. glutamicum* wild type.** An OmniLog System from Biolog (Hayward CA, USA) was used to perform phenotypic microarrays with wild type *C. glutamicum* ATCC 13032 as well as the *malR* deletion strain. The experiments (PM1, PM2A, PM4, PM9, PM10, PM11, PM12, PM13) were conducted like described in the protocol of the manufacturer (Bochner BR, Gadzinski P, Panomitros E. 2001. Genome Res 11:1246–1255.). The wild type strain is displayed in red, whereas the *malR* deficient mutant is green. Yellow means a similar behavior.
