## Supplementary material for "The MarR-type regulator MalR is involved in stress-responsive cell envelope remodeling in *Corynebacterium glutamicum*": Table S4

### Supporting Information

#### Table S4

**Table S4: Antibiotics affecting the growth of wild type cells or Δ*malR* cells during phenotype microarrays (BioLog).** For evaluation of the sensitivity of the wild type ATCC 13032 or the *malR* deletion mutant, the BioLog plates PM11, PM12 PM13 were used. The table shows antibiotics were both strains show differences in the metabolic activity.

| Antibiotic | Better Growth | Substance class |
|---|---|---|
| Gentamicin | Δ*malR* | Aminoglycoside |
| Amikacin | Δ*malR* | Aminoglycoside |
| Lincomycin | wt | Lincosamide |
| Erythromycin | Δ*malR* | Glycoside |
| Chlortetracyclin | wt | Tetracycline |
| Demeclocycline | wt | Tetracycline |
| Tetracyclin | wt | Tetracycline |
| Penimepicyclin | wt | Tetracycline |
| Cefazolin | wt | β-Lactame; Cephalosporine |
| Cephalothin | wt | β-Lactame; Cephalosporine |
| Cefuroxime | wt | β-Lactame; Cephalosporine |
| Amoxicillin | wt | β-Lactame; Penicilline |
