## Supplementary material for "The MarR-type regulator MalR is involved in stress-responsive cell envelope remodeling in *Corynebacterium glutamicum*": Table S3

**Supporting Information**

**Tables S3 A, B, C**

Table S3 A: Oligonucleotides used in this study

| **Number** | | **Sequence (5’-3’)** |
| --- | --- | --- |
| 1 | | AGGTCGACTCTAGAGGATCCTCGAACGGAAATTACTTGGCAATAC |
| 2 | | CTTCTCGAACTGTGGGTGGGACCAGCTAGCAGAACCGCTGTCGGTCT |
| 3 | | GCTAGCTGGTCCCACCCACAGTTCGAGAAGTAACAGTTTTCTCCATCTCAACTCC |
| 4 | | AAACGACGGCCAGTGAATTCACAAGTCCTAGTGGGACGG |
| 5 | | AGGTCGACTCTAGAGGATCCCCGTCCATTCTTTACCAACTGT |
| 6 | | CCCATCCACTAAACTTAAACAAGCGGTATTGCCAAGTAATTTCC |
| 7 | | TGTTTAAGTTTAGTGGATGGGCAGTTTTCTCCATCTCAACTCCG |
| 8 | | AAACGACGGCCAGTGAATTCGGCACAAGTCCTAGTGGGAC |
| 9 | | TTTAAGAAGGAGATATACATATGCTGGCAGGCATGC |
| 10 | | AAATACAGGTTCTCGCTAGCAGAACCGCTGTCGGTCTC |
| 11 | | AAGCTTGCATGCCTGCAGAAGGAGGAGTCGTATGCCTAATTTAAACGCTGAGGAG |
| 12 | | AAACGACGGCCAGTGAATTCTTAAGAACCGCTGTCGGTCTC |
| 13 | | GCCAAGGTAAGTTGTACTTTTTCTG |
| 14 | | GTAGTTGACGCGGGTGAC |
| 15 | | TAACAATCCCAACTCGAAGCAC |
| 16 | | TGACAATCCTCTCCACGAAGC |
| 17 | | GAGTGTGAGAAACATGGGACGTG |
| 18 | | ACAGGTTACCAGCCAAAGTG |
| 19 | | TCGATACCCAGAAAGAATTGCATTTG |
| 20 | | ATCTCCCTGGATAGTGATCTTGC |
| 21 | | CCGTCCAGAACTAGGACTATTTG |
| 22 | | TAATCCCAACAATCGCTTATGACG |
| 23 | | TTGAAATCGTGATCGCCTGTTATTG |
| 24 | | AGCCATGTTTGCTTCTCCTTTTC |
| 25 | | TCAGGTTTACTACCGACTGTAGTAG |
| 26 | | TTTGAAAGTAGAAATTACTGGTAGTGGATTA |
| 27 | | TTTACCTTCCTCTACCTAATCTCCC |
| 28 | | ATCTCAGTAGCGTTGCCTCC |
| 29 | | TTTCTCATTTCGCCACCCCC |
| 30 | | CTAAGATGAGTATTTAAGCCCTGTTTAT |
| 31 | | ACGCCCGTATCGTTTCGC |
| 32 | | ATGTACTTGTTTAAAACAATAGTTGTCAATG |
| 33 | | GCACCCATGGTTAGCGTACT |
| 34 | | TTCTACCGGGGGTAGTAGCG |
| 35 | | TGTTGCTTCCGCTGACC |
| 36 | | ACCTTCTATATAAACCTTTTATGAGGGAAATG |
| 37 | | CTGGACCGAGCTCAATGC |
| 38 | | TGAAAAATTGTTTTAATATGCAACACAACTA |
| 39 | | TTGTAGGCGTTGGACACTG |
| 40 | | AGATTGAAATGATTATGGGTAGGAAAC |
| 41 | | AAGAGCCGTGATGTTAACAAATG |
| 42 | | TAGTTAGGTTACACTAATGGTGTCC |
| 43 | | GGAACAGCACAGAATTAAGGC |
| 44 | | GAAAAAGCAATAATTTGGACAGAAAAAG |
| 45 | | AAACCACCCCTGTACAAAATTAGC |
| 46 | | AAAAACACTAGCTAAATCTGTAGCTCAAC |
| 47 | | ATGTTTCTTACTTTAAGCGCAGTTAATTG |
| 48 | | GGATTTAAATTTTGTATTGGAAAGCTAATAATT |
| 49 | | CATAGGGGTATAGCCTTGAG |
| 50 | | CAGTGTGCGCAGGTCATGCC |
| 51 | | GTATGGGTTGAGATTCCGACAG |
| 52 | | TCTTCCCTATCACCTCCAGTT |
| 53 | | GGAGCTTTCGCTGACTATCTT |
| 54 | | AATTCTCCTGCGTCGTCTTT |
| 55 | | GACGTGGATCGTGTATGGAATTA |
| 56 | | CAGCCTTCTCAAGGTGGATAAA |
| 57 | | ACTGTGGATCGACATTCCTTTC |
| 58 | | CCGCGTCTAAGTCCCTTTAATC |
| 59 | | ATGACTCGATCCCAGGACTAT |
| 60 | | TGAATGCGGTATGAGCTAAGG |
| 61 | | GCCCAAACACCACCGATATT |
| 62 | | CATCAGTCAAGCAGGTCTGAAC |
| 63 | | CATCTTCCATCCACAGACCTAAT |
| 64 | | GAAGTCGGACACGATGTAGAAG |
| 65 | | ATCTCGTGCTTGCTGTGATTA |
| 66 | | CTGGGTGAATCCTAAAGACCTG |
| 67 | | CTGATGGGTCAGTCGTAGTTTC |
| 68 | | CCGAATAGAGCCAGGAACAAT |
| 69 | | CTCGACAACGTGAAGCTGTTA |
| 70 | | CAAGCCAGGTCTGTGTGATT |
| 71 | | GCTGTAGGTAAGGGCTTTGATA |
| 72 | | TAGCCAACAGCCTGGTAATC |
| 73 | | GAGCTTCAGAAACTTGCAACAG |
| 74 | | CATTGAGGGCGAGGATGATT |
| 75 | | AGCGAAAGTTCCCGAATCTG |
| 76 | | CGGACGGTCAGTCTTGTTATG |
| 77 | | CTCGACACCGCTGAAGATATG |
| 78 | | CAGTAAGCAGCACTCCGATT |
| 79 | | GGGTCCACAGCTCACTATTT |
| 80 | AGCGGTTGGCATACGAATA | |
| 81 | | AGCAATCAAGGAAGATCCAGAG |
| 82 | | GATATCGCCAAGGCCAAGAA |
| 83 | | CCGGTTCTTCTGCATCTTCT |
| 84 | | GGTTACATCGTCGAAGTCCTTA |
| 85 | | CAGACTCACAACAACGTCAAAC |
| 86 | | CTAGTTCGTGGCCAACTTCA |
| 87 | | TTCCTCACAGATCGCTTTCG |
| 88 | | GAGCAGGTATGGAGCAACTT |

**Table S3 B: Construction of plasmids used in this study.** Numbers represent oligonucleotide pairs used for PCR (see Table S3 A). The restriction enzymes were used for linearization of the vectors and plasmids were assembled using Gibson assembly. Sequencing for verification of the chromosomal modifications was conducted using primers 13 + 14.

| **Plasmid** | **Template** | **Primers** | **Vector** | **Restriction Enzymes** |
| --- | --- | --- | --- | --- |
| pEKEx2-*malR* | *C. glutamicum* chromosome | 11 + 12 | pEKEx2 | *PstI *EcoRI |
| pK19-*malR*-C-*strep* | *C. glutamicum* chromosome | 1 + 2; 3 + 4 | pK19*mobsacB* | *BamHI *EcoRI |
| pK19-Δ*malR* | *C. glutamicum* chromosome | 5 + 6; 7 + 8 | pK19*mobsacB* | *BamHI *EcoRI |
| pET24b-*malR*-C-*strep* | *C. glutamicum* chromosome | 9 + 10 | pET24b | *NheI *NdeI |

**Table S3 C: Amplification of EMSAs DNA probes.** Indicated are the oligonucleotide pairs used for PCR amplification of the 100 bp fragments. The fragments cover the maximum peak position determined by ChAP-Seq and lay inside the promoter regions of each gene.

| **Fragment** | **Gene** | **Oligonucleotides (Table S1)** |
| --- | --- | --- |
| 1 | cg0111 | 15 + 16 |
| 2 | cg0120 | 17 + 18 |
| 3 | cg0423 | 19 + 20 |
| 4 | cg0438 | 21 + 22 |
| 5 | cg1577 | 23 + 24 |
| 6 | cg1905 | 25 + 26 |
| 7 | cg1909 | 27 + 28 |
| 8 | cg1911 | 29 + 30 |
| 9 | cg1929 | 31 + 32 |
| 10 | cg2033 | 33 + 34 |
| 11 | cg2500 | 35 + 36 |
| 12 | cg2904 | 37 + 38 |
| 13 | cg2910 | 39 + 40 |
| 14 | cg2962 | 41 + 42 |
| 15 | cg3315 | 43 + 44 |
| 16 | cg3335 | 45 + 46 |
| 17 | cg3343 (positive control) | 47 + 48 |
| 18 | cg3402 (negative control) | 49 + 50 |

**Table S3 D: Oligonucleotide combinations for qRT-PCR analysis to verify microarray data.** Indicated are the oligonucleotide pairs used for qRT-PCR. Each pair comprises approximately 100 bp fragments and shares similar features regarding GC-content and melting temperature.

| **Fragment** | **Gene** | **Oligonucleotides (Table S1)** |
| --- | --- | --- |
| 1 | cg0111 | 51 + 52 |
| 2 | cg0120 | 53 + 54 |
| 3 | cg0422 | 55 + 56 |
| 4 | cg0423 | 57 + 58 |
| 5 | cg0438 | 59 + 60 |
| 6 | cg1577 | 61 + 62 |
| 7 | cg1905 | 63 + 64 |
| 8 | cg1909 | 65 + 66 |
| 9 | cg1911 | 67 + 68 |
| 10 | cg1929 | 69 + 70 |
| 11 | cg2033 | 71 + 72 |
| 12 | cg2500 | 73 + 74 |
| 13 | cg2904 | 75 + 76 |
| 14 | cg2910 | 77 + 78 |
| 15 | cg3315 | 79 + 80 |
| 16 | cg3335 | 81 + 82 |
| 17 | cg3343 | 83 + 84 |
| 18 | cg3206 | 85 + 86 |
| 19 | cg2900 | 87 + 88 |
