## Supplementary material for "The MarR-type regulator MalR is involved in stress-responsive cell envelope remodeling in *Corynebacterium glutamicum*": SI Methods

### Supporting Information – Methods & Table S3

#### Methods

##### Additional information to growth conditions

For the cultivation of *C. glutamicum,* a first pre-culture was inoculated with single colonies from agar plates either directly after transformation or after a streak-out of glycerol cultures. These pre-cultures were conducted in 4.5 ml BHI medium in test tubes or – depending on the purpose - in 1 ml BHI medium in 96-well deep well plates (DWPs) at 30°C for 8 h. Afterwards cells were used for inoculating a second overnight pre-culture in CGXII medium containing 2% (w/v) glucose. This CGXII culture was used to inoculate a main-culture in the same medium to an $OD_{600}$ of 1.

##### Scanning and transmission electron microscopy.

For scanning electron microscopy (SEM), bacteria were fixed with 3% (vol/vol) glutaraldehyde (Agar Scientific, Wetzlar, Germany) in PBS for at least 4 h, washed in 0.1 M Soerensen's phosphate buffer (Merck, Darmstadt, Germany) for 15 min, and dehydrated by incubating consecutively in an ascending acetone series (30%, 50%, 70%, 90%, and 100%) for 10 min each and the last step thrice. The samples were critical point dried in liquid $CO_2$ and then sputter coated (Sputter Coater EM SCD500; Leica, Wetzlar, Germany) with a 10-nm gold/palladium layer. Samples were analyzed using an environmental scanning electron microscope (ESEM XL 30 FEG, FEI, Philips, Eindhoven, Netherlands) with a 10-kV acceleration voltage in a high-vacuum environment.

Transmission electron microscopy.

In preparation for transmission electron microscopy (TEM) analysis, bacteria were fixed with 3% (vol/ vol) glutaraldehyde (Agar Scientific, Wetzlar, Germany) in PBS for at least 4 h, washed in 0.1 M Soerensen's phosphate buffer (Merck, Darmstadt, Germany), and postfixed in 1% OsO4 in 17% sucrose buffer. After fixation, bacteria were embedded in 2.5% agarose (Sigma, Steinheim, Germany), then rinsed in 17% sucrose buffer and deionized water and dehydrated by ascending ethanol series (30, 50, 70, 90 and 100%) for 10 min each. Last step was repeated 3 times. Dehydrated specimens were incubated in propylene oxide (Serva, Heidelberg, Germany) for 30 min, in a mixture of Epon resin (Serva, Heidelberg, Germany) and propylene oxide (1:1) for 1h and finally in pure Epon for 1h. Samples were embedded in pure Epon. Epon polymerization was performed at 90°C for 2h. Ultrathin sections (70-100 nm) were cut by ultramicrotome (Reichert Ultracut S, Leica, Wetzlar, Germany) and picked up on Cu/Rh grids (HR23 Maxtaform, Plano, Wetzlar, Germany). Contrast was enhanced by staining with 0.5% uranyl acetate and 1% lead citrate (both EMS, Munich, Germany). Samples were viewed at an acceleration voltage of 60 kV using a Zeiss Leo 906 (Carl Zeiss, Oberkochen, Germany) transmission electron microscope.
